# A Computational Foundation Toward Targeting the ELMO1/DOCK2 Complex

**DOI:** 10.64898/2026.07.27.740996

**Authors:** Soumita Das, Anastasia Ignashkina, Hossam Hammouda

**Affiliations:** Department of Biomedical and Nutritional Sciences, University of Massachusetts Lowell, Lowell, MA, USA

**Keywords:** ELMO1, DOCK2. FEL, MMGBSA

## Abstract

Engulfment and Cell Motility protein 1 (ELMO1) regulates cell migration, phagocytosis, and cytoskeletal remodeling, positioning it as a compelling therapeutic target across kidney diseases, oncology, enteric infections and inflammation. Despite this potential, no approved therapeutics or clinically validated small-molecule modulators of ELMO1 currently exist. ELMO1 functions by forming a complex with DOCK180 (or DOCK2) to activate the small GTPase Rac1, and the recent structural resolution of the ELMO1/DOCK2 complex now provides an opportunity to target this protein–protein interface directly. Here, we present the first investigation into the druggability of the ELMO1/DOCK2 complex and report the initial virtual screening to identify small-molecule inhibitors of this interaction. Molecular dynamics (MD) and free energy level (FEL) studies were carried out to validate the potential of the predicted hits. This work establishes a computational framework for the development of the first generation of ELMO1-targeted therapeutics. In addition to demonstrating the drugability of ELMO1, this work introduces two open-source Python tools for the rapid analysis and visualization of protein– protein interaction and ligand–protein MD trajectories from DESMOND output files. These tools are designed to be broadly accessible, offering practical utility to the wider DESMOND user community.

## 1. Introduction

Engulfment Cell Motility Protein 1 (ELMO1) is a cytosolic protein best known for its role in regulating actin organization and the engulfment of apoptotic cells and enteric bacteria ^1-3^. Previously, we have shown that ELMO1 binds the pattern-recognition receptor BAI1 (Brain angiogenesis Inhibitor 1), which recognizes Gram-negative bacteria via lipopolysaccharide (LPS) ^3^. In bacterial infection, ELMO1 acts as a microbial sensor by sensing and generating inflammation against pathogenic bacteria compared to non-pathogenic bacteria through interaction with the WxxxE effectors of enteric bacteria ^4^.The engulfment is triggered by the BAI1-mediated activation of Rac through an ELMO/Dock-dependent mechanism ^3^. ELMO1, after activation of Rac1, induces the expression of pro-inflammatory cytokines. The dedicator of cytokinesis (DOCK) protein family interacts with ELMO1 and acts as a GEF for Rho GTPases ^5^. Of the 11 DOCK proteins, only DOCK 1-5 bind the ELMO domain of ELMO1. (PMID: 12134158). Recent cryo-EM findings showed that DOCK5, ELMO1, RhoG, and Rac1 are aligned on a single plane and symmetrically flattened ^6^. Physiologically, the DOCK-ELMO complex is important and has been shown in 1) T cells and neutrophils that migrate towards the site of infection and inflammation ^7^; 2) CD11c–DOCK2–ELMO1 interaction in neutrophils activates Rac for ROS formation ^8^. The HIV-1 virulence factor Nef activates Rac by binding the DOCK2–ELMO1 complex, and this interaction is likely linked to inhibition of chemotaxis and activation of T cells ^9^.

Mechanistically, the ELMO1–DOCK2 complex is held in an autoinhibited “locked” state at rest ^10^ (Figure 1) where ELMO1 folds back onto DOCK2 leading to the occlusion of the catalytic DHR2 domain and leaving Rac1 in its inactive, GDP-bound form ^11^. This autoinhibition is relieved by upstream activating cues such as engagement of chemokine receptors (e.g., CXCR) by chemokines such as CXCL, which triggers RhoG activation, and RhoG binding to ELMO1 opens the complex ^12-14^. TAM kinase-mediated phosphorylation of complex components further facilitates this unlocking, exposing the previously hidden DHR2 domain. Once exposed, the DOCK2 DHR2 domain functions as the catalytic GEF module where it binds to Rac1-GDP and catalyzes the release of GDP, which is followed by GTP loading ^15-17^. This process leads to the generation of active Rac1-GTP. Active Rac1 then recruits downstream effectors at the leading edge of the cell, including the WAVE complex, PAK1, and other effectors, driving actin filament polymerization and lamellipodia formation. This actin remodeling underlies the directional cell movement, chemotaxis, and phagocytic activity associated with ELMO1–DOCK2–Rac1 signaling ^18^. The crystal structure of DOCK2 and ELMO1 has shown that they mutually relieve their autoinhibition for the activation of Rac1 for lymphocyte chemotaxis ^19^. Stevenson et al. have also shown that ELMO1 controls DOCK2 levels and DOCK2-dependent T cell migration ^20^.

**Figure 1.**
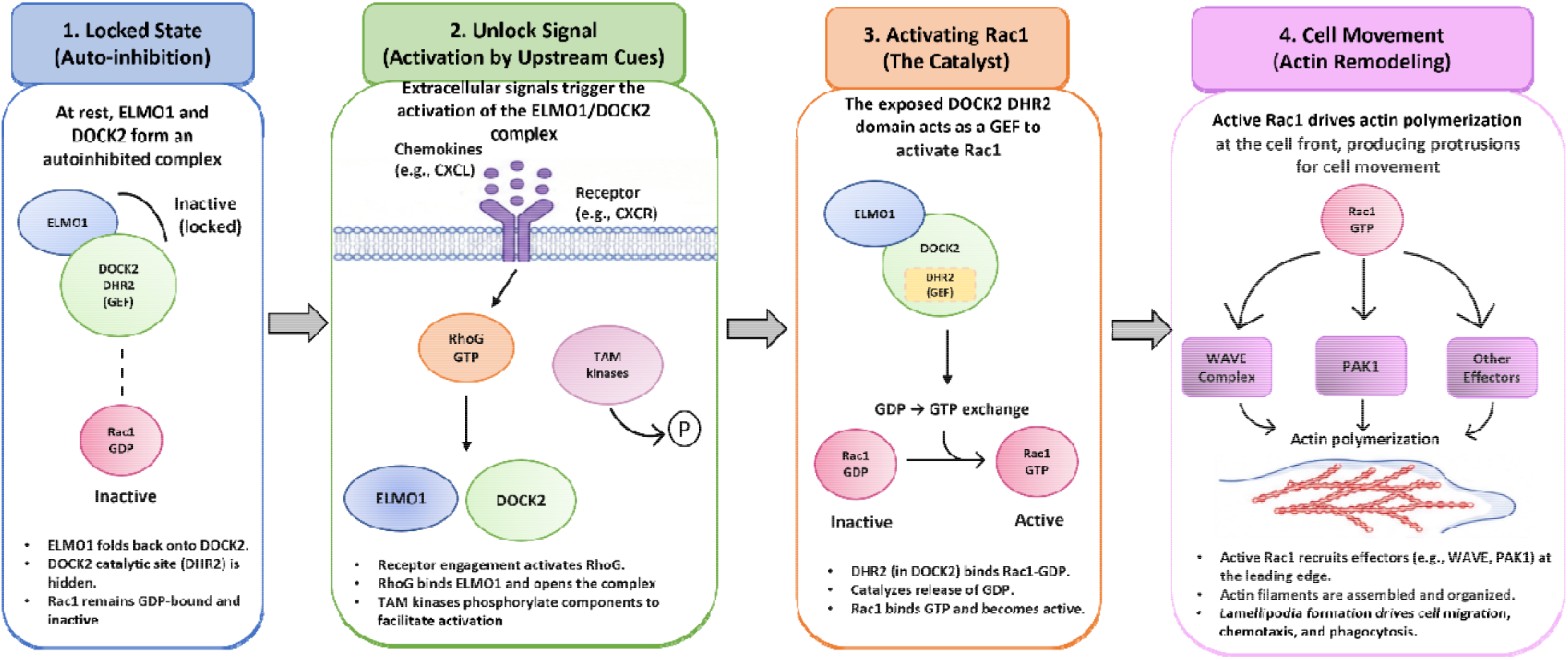
Signaling mechanism of ELMO1–DOCK2-mediated Rac1 activation and downstream cytoskeletal remodeling. The ELMO1–DOCK2 complex regulates Rac1 activity through a stepwise, four-part mechanism. (1) Locked state: auto-inhibition. (2) Unlock signal: activation by upstream cues. (3) Activating Rac1: the catalyst. (4) Cell movement: actin remodeling. (**Other functions of active Rac1 in controlling inflammation and the related papers have not been shown here**)

Despite this well-defined signaling architecture, the ELMO1–DOCK2 complex remains pharmacologically unexplored, as no small molecules have been reported to modulate this protein-protein interaction (PPI). Herein, we establish a computational framework for targeting the ELMO1/DOCK2 complex, using the autoinhibited conformation as the structural basis for interface characterization and pocket identification. This work assesses the druggability of the ELMO1/DOCK2 interface and reports the first virtual screening campaign directed at identifying small-molecule inhibitors of this PPI. Predicted hits were subsequently subjected to molecular dynamics (MD) simulations and Free Energy Landscape (FEL) analysis to evaluate binding pose stability and validate the potential of the predicted hits in modulating the PPI. Collectively, this work establishes a computational pipeline for the development of a first-generation of ELMO1-targeted therapeutics. Beyond demonstrating the druggability potential of ELMO1 as a therapeutic target, this study introduces two Python scripts for the rapid analysis and visualization of PPI and ligand-protein MD trajectories. Built around DESMOND output files, these tools are intended to be broadly useful to the wide community of DESMOND users engaged in PPI and small-molecule MD analysis.

## 2. Methods

### 2.1. ELMO1/DOCK2 interaction analysis

A 100 ns MD simulation was carried out to investigate the interaction of ELMO1 with DOCK2 (PDB ID: 6TGB ^11^) using Desmond at default settings. The MD simulation yielded 1,001 trajectory frames, which were sampled at uniform intervals. Simulation output was stored in Schrödinger Event Analysis Format (.eaf) files, which encode per-frame interaction data including residue-level contact events, RMSD and RMSF series, secondary structure assignments, and protein–protein interaction (PPI) summaries.

The resulting files were analyzed using our custom-built Python script “PPI.py” which has been provided in the supplementary files. The frequency of ELMO1/DOCK2 contact was computed as the number of frames in which the contact was observed divided by the total number of trajectory frames. The contact between the two protein chains was classified by interaction type into backbone–backbone hydrogen bonds, sidechain–sidechain hydrogen bonds, mixed backbone–sidechain hydrogen bonds, salt bridges, π-π stacking, and π–cation interactions. The Python script employs the inter-chain contact frequency to classify the observed interaction residues into hotspot residues (contact frequency of more than 70%, interface residues (contact frequency of 30-70%), and peripheral residues (contact frequency of less than 30%.

Conformational sampling was assessed through two-dimensional free energy landscapes (FEL) constructed using backbone RMSD and radius of gyration as collective variables. Normalized probability densities were computed over a 60-bin grid, Gaussian-smoothed (σ = 1.5 bins), and converted to free energy which was referenced to zero at the global free energy minimum. Backbone flexibility was assessed via per-residue Cα RMSF profiles. For visualization, a companion script (Visualization.py, Supplementary File) generated a multi-panel publication figure depicting ELMO1 primary (residues 3–128) and secondary (C-terminal, residues 1341–1525) binding contact frequencies, the DOCK2 complementary interface, a two-dimensional binding schematic annotating dominant pairwise interactions, and an interaction-type distribution summary. The script renders all figures at 300 dpi using Matplotlib and saved in both PNG and PDF formats.

### 2.2. Virtual screening

The ELMO1-identified binding site was utilized to generate a grid for virtual screening. Structure-based virtual screening was performed using the Maestro Schrödinger suite where a focused chemical library of 15,000 compounds was assembled from the Enamine Diversity Discovery Set (DDS) and screened against the identified ELMO1 binding pocket (PDB ID: 6TGB). The 15,000 Enamine DDS subset compounds were prepared using the LigPrep module ^21^, which generated tautomeric and ionization states at pH 7.0 ± 2.0, optimized geometries, and produced low-energy 3D conformers for docking input. The virtual screening campaign consisted of three stages. The first stage of the virtual screening campaign involved high-throughput virtual screening (HTVS), which was performed on all 15,000 compounds using a simplified scoring function to rapidly eliminate non-binders. The top 10% ranking compounds progressed to standard precision (SP) docking for more accurate pose prediction and scoring. Finally, the top 10% predicted candidates underwent extra precision (XP) docking with enhanced sampling and a more rigorous scoring function to identify the most promising hits for downstream experimental validation.

### 2.3. Molecular Dynamics (MD)

MD simulations were performed using the Desmond simulation package (Schrödinger, LLC) for the complexes of compounds **SH 1–5** with ELMO1. Each simulation system was constructed within an orthorhombic periodic boundary box, solvated with explicit SPC water molecules, and neutralized with counterions to achieve a physiologically relevant ionic environment ^22^. All simulations were conducted at 300 K and 1 bar pressure (NPT ensemble) for a total production run of 100 ns, with a neighbor list update interval and positional relaxation time of 1 ps. Intermolecular interactions were parameterized using the OPLS 2005 force field. Long-range electrostatic interactions were computed using the Particle Mesh Ewald (PME) method, and a cutoff radius of 9.0 Å was applied to non-bonded short-range interactions. Prior to production runs, each system underwent a six-stage relaxation protocol implemented within Desmond to progressively relieve steric clashes, equilibrate solvent density, and stabilize temperature and pressure before data collection. Post-simulation analysis, including trajectory processing and free energy landscape (FEL) construction, was performed using a custom in-house Python script (MD.py). MMGBSA calculations were carried out using thermal_mmgbsa.py script.

## 3. Results and discussion

### 3.1. ELMO1/DOCK2 interaction analysis

A 100 ns MD simulation was carried out to explore the interaction of ELMO1 with DOCK2, which revealed that the complex undergoes rapid initial structural rearrangement within the first ∼10 ns, after which the system enters a phase of dynamic equilibration that is sustained through the remainder of the trajectory (Figure 2A). The near-parallel evolution of backbone and sidechain RMSD profiles indicates that the observed conformational changes are likely due to collective domain motion rather than localized sidechain reorganization, as supported by the relatively stable plateau of the mean RMSD of 10.1 Å. Subsequent analysis of the protein-protein Interface led to the identification of the key residues that are predicted to be crucial to the stability of the ELMO1/DOCK2 complex. On the ELMO1 side, GLU-693 and ASP-685 emerged as the dominant hotspot residues, where they were calculated to maintain contact frequencies of 98.6% and 91.1%, respectively, throughout the whole simulation. CYS-726 and GLU-535 also exceeded the 70% hotspot threshold, while a broader set of residues, including PRO-712, GLU-702, VAL-723, TYR-724, and ASP-725, contributed to a more extended interface zone.

**Figure 2.**
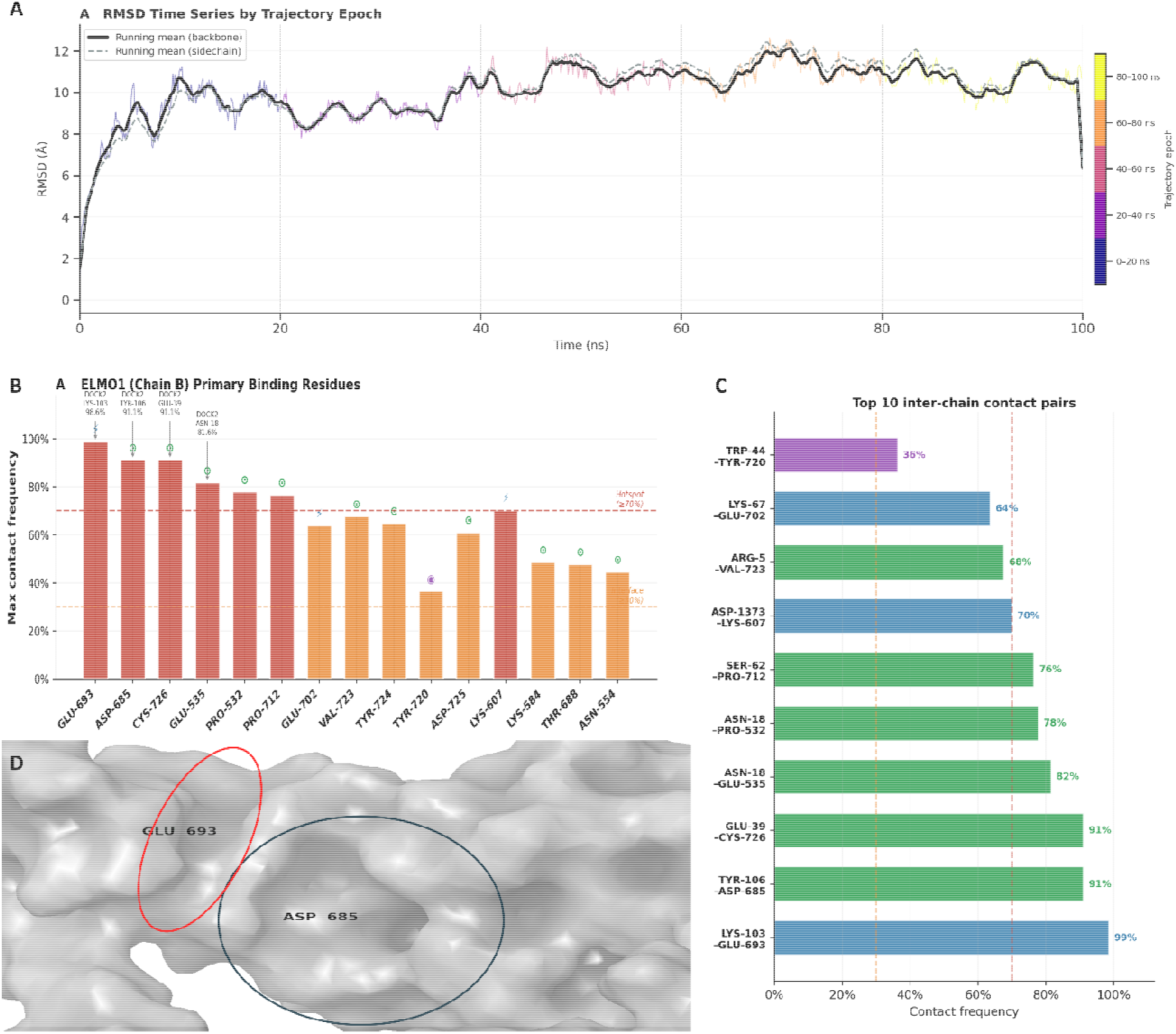
Molecular dynamics simulation of the ELMO1–DOCK2 protein–protein interface. (A) Backbone (solid black) and sidechain (dashed grey) RMSD time series over the 100 ns trajectory. (B) Maximum inter-chain contact frequencies for ELMO1 primary binding residues (chain B), with hotspot (≥ 70%, red dashed) and interface (≥ 30%, orange dashed) thresholds indicated. (C) Ranked bar chart of the top 10 inter-chain residue–residue contact pairs, color-coded by interaction type (blue, salt bridge; green, hydrogen bond; purple, π–π stacking), with contact frequencies labeled. (D) Molecular surface representation of the ELMO1 binding groove.

Subsequently, the binding interface of ELMO1 was visualized (Figure 2D) at 20ns of the MD simulation (most stable RMSD), where a possible binding pocket was observed featuring two sub-pockets centered on GLU-693 and ASP-685. This binding site corresponded to the highest-frequency contact hotspots identified in the quantitative analysis of the ELMO1/DOCK2. These structural observations provide a rational basis for targeting the ELMO1/DOCK2 interface by conducting a virtual screening campaign capable of targeting the two key residues GLU-693 and ASP-685.

### 3.2. Virtual screening

Subsequently, structure-based virtual screening identified 15 potential hits with favorable predicted binding affinities to the ELMO1 pocket. Next, visual analysis of the predicted hits identified five compounds that occupied the two sub-pockets of the predicted binding site and established interactions with ASP-685 and GLU-693 of ELMO1. These compounds (Z1011345768, Z505786020, Z4872634012, Z1277268301 and Z2058591482) were subsequently renamed as **SH 1-5**, respectively. The docked poses of **SH 1-5** in complex with ELMO1, as well as the unoccupied binding pocket, are illustrated in Figure 3.

**Figure 3.**
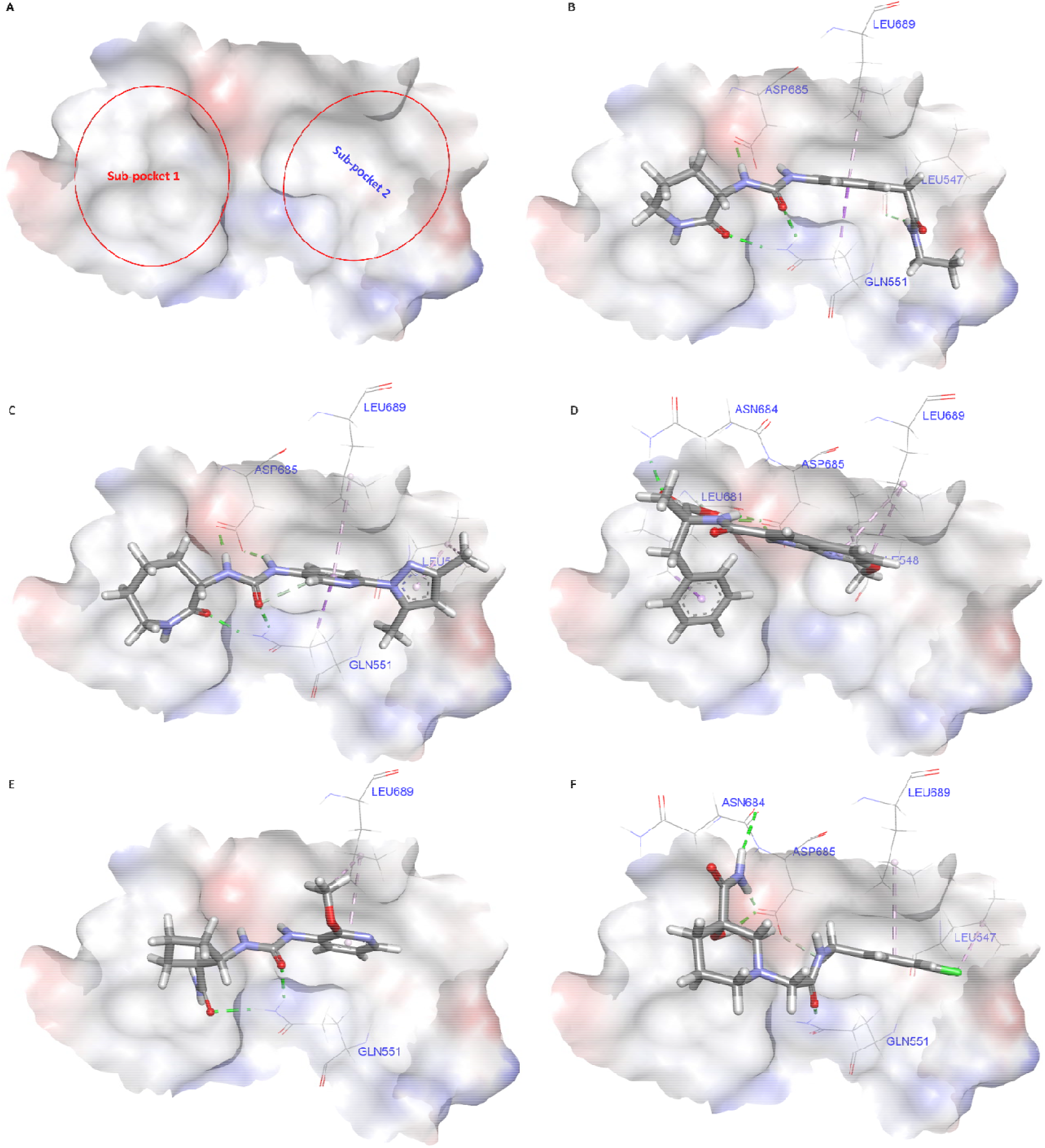
Structural analysis of the ELMO1 binding pocket and docking poses of top virtual screening hits SH1–SH5. (A) Surface representation of the predicted unoccupied ELMO1 binding pocket (PDB ID: 6TGB) rendered by electrostatic potential (red, negative; blue, positive; white, neutral), illustrating the two distinct sub-pockets targeted in virtual screening: Sub-pocket 1 (left, larger hydrophobic cavity) and Sub-pocket 2 (right, polar sub-region). (B–F) Docking poses of the five top-ranked hits **SH1** (B), **SH2** (C), **SH3** (D), **SH4** (E), and **SH5** (F) shown as stick models within the ELMO1 electrostatic surface. Key interacting residues are labeled. Green dashed lines indicate hydrogen bonds; magenta dashed lines indicate hydrophobic contacts.

The predicted ELMO1 pocket (Figure 3A) was presented as a “bilobal” cavity comprising a larger, predominantly hydrophobic Sub-pocket 1 and a smaller, polar Sub-pocket 2 enriched in hydrogen-bond donors and acceptors contributed by ASN-684, ASP-685, and GLN-551. Given that both ASP 685 of the polar sub-pocket and GLU-693 of the hydrophobic sub-pocket were involved in persistent ELMO1/DOCK2 contact, dual sub-pocket occupancy by a small molecule is anticipated to be a key determinant of successful PPI modulation.

**SH1, SH2**, and **SH3** (Figures 3B and 3C) all adopted an extended, linear binding pose, which was calculated to occupy both sub-pockets simultaneously. In the hydrophobic sub-pocket 1, both **SH1** and **SH2** established hydrophobic interactions with LEU689 and LEU547 of ELMO1. Meanwhile, both compounds established hydrogen bond interactions with ASP685 and GLN551 of the polar sub-pocket which indicates favorable electrostatic complementarity. Meanwhile, while **SH 4-5** managed to partially occupy both sub-pockets, they only fully occupied one, indicating that they might possess weaker inhibitory activity. Subsequently, molecular dynamics studies were carried out to validate the results of the virtual screening and evaluate the conformational behavior of the **SH 1-5** complexes within a flexible, biologically relevant environment.

### 3.3. Molecular dynamics

Five molecular dynamic simulations were carried out to validate the predicted binding modes of hit compounds **SH 1-5** with ELMO1. The stability of the ligand–ELMO1 predicted complexes was first evaluated by observing the root-mean-square deviation (RMSD) of the complexes. Among the five predicted hits, **SH1–4** (Figure 4) displayed broadly acceptable stability profiles with enhanced stability when compared to the unbound protein complex (Figure 2A). Meanwhile, **SH5** exhibited anomalous behavior, with a highly fluctuating RMSD, indicating that it is a false positive (Figure 4E).

**Figure 4.**
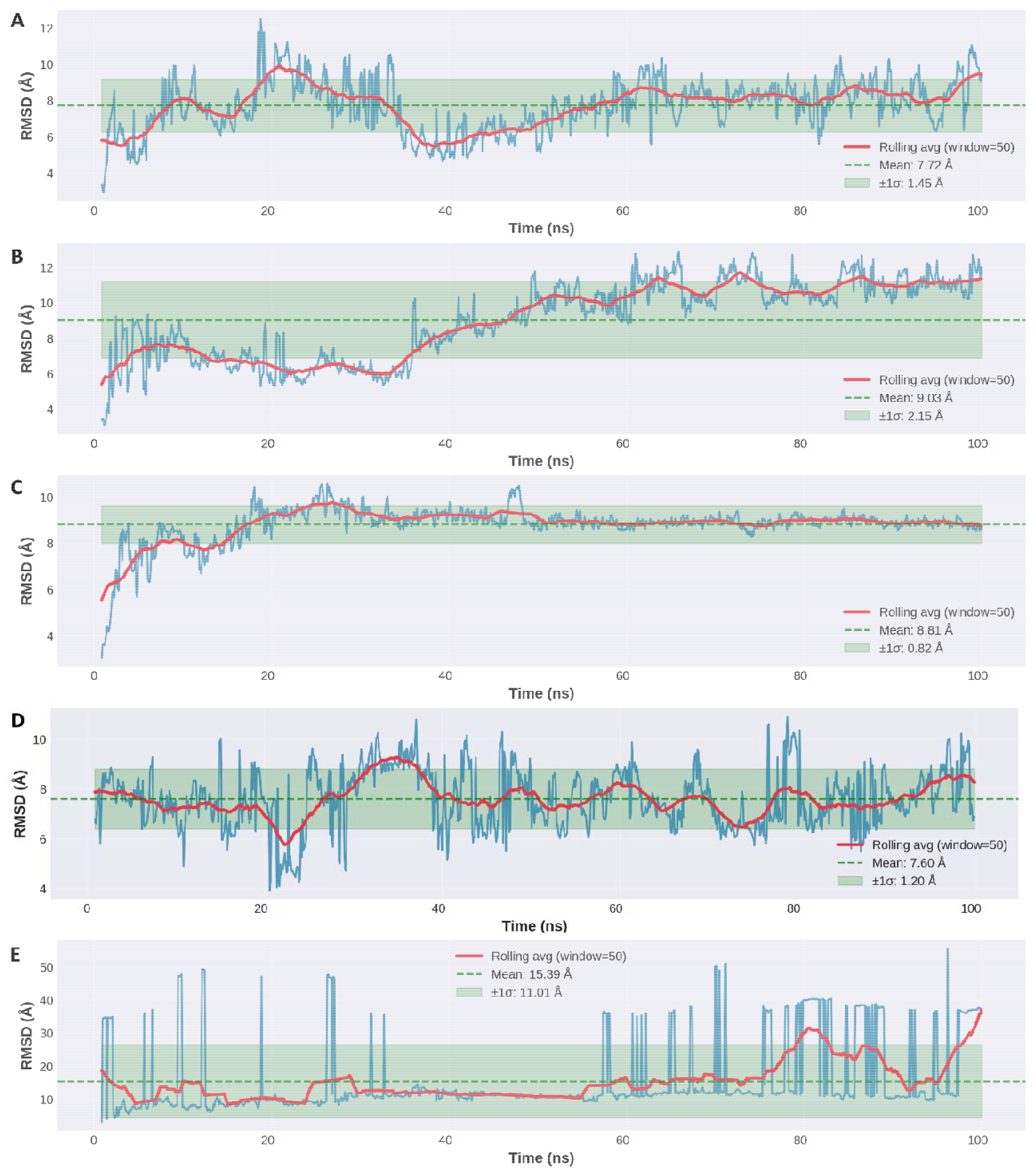
RMSD of ELMO1 in complex with SH 1-5 over 100 ns molecular dynamics simulations. RMSD of the ELMO1 protein backbone (Cα atoms) relative to the initial docked conformation is shown for each of the five hit complexes: (A) **SH1**/ELMO1, (B) **SH2**/ELMO1, (C) **SH3**/ELMO1, (D) **SH4**/ELMO1, and (E) **SH5**/ELMO1. Raw RMSD values are plotted in blue; the red line represents the rolling average while the green dashed line indicates the mean RMSD over the full trajectory, and the green shaded region denotes the ±1 standard deviation (σ) envelope.

The **SH1**/ELMO1 complex (Figure 4A) exhibited the lowest RMSD, with a mean backbone RMSD of 7.72 ± 1.45 Å over the 100 ns MD simulation. **SH1**/ELMO1 complex exhibited an initial equilibration phase over the first 20 ns, which was accompanied by moderate fluctuations due to local structural reorganization, which equilibrated to the baseline RMSD of approximately 6–7 Å before gradually stabilizing at approximately 8 Å through the remainder of the simulation. The relatively contained standard deviation indicates that, despite the early perturbation, the **SH1**/ELMO1 complex reached a stable binding mode and maintained it for most of the production run, supporting the viability of SH1 as a lead candidate.

The **SH2**/ELMO1 complex (Figure 4B) exhibited the highest mean RMSD of 9.03 ± 2.15 Å among the five systems and the largest standard deviation among the four nominally stable complexes. Rather than converging to a stable plateau, the trajectory displayed a progressive upward drift from approximately 6 Å at the outset to 10–11 Å by 60 ns, with elevated fluctuations persisting through the end of the simulation. The fluctuations observed with the **SH2**/ELMO1 complex, along with the observed inability to converge, indicate that the complex is not stable enough and either needs further optimization or warrants further investigation to assess its predicted stability.

In contrast, the **SH3**/ELMO1 complex (Figure 4C) demonstrated the most favorable stability profile among the five predicted hit compounds examined. The **SH3**/ELMO1 complex exhibited a mean RMSD of 8.81 Å and a standard deviation of only 0.82 Å, which was the lowest of all five systems. Following an initial rise from approximately 3 Å during the equilibration phase, the RMSD plateaued at approximately 8.8–9.0 Å by 30 ns and remained essentially invariant for the remaining 70 ns of the trajectory. This rapid convergence and minimal fluctuations around a stable mean are hallmarks of a well-accommodated ligand– protein interaction and indicate that SH3 adopts a single, thermodynamically preferred binding mode within the ELMO1 pocket that is resistant to conformational drift.

The **SH4**/ELMO1 complex (Figure 4D) achieved the lowest mean RMSD across the series at 7.60 ± 1.20 Å, which reflects compact and contained backbone dynamics throughout the simulation. The trajectory oscillated around the mean with periodic, moderate fluctuations but without any sustained upward drift or destabilizing events. Lastly, the SH5/ELMO1 complex (Figure 4E) exhibited a mean RMSD of 15.39 Å and a standard deviation of 11.01 Å, where the trajectory displayed large-amplitude, episodic spikes reaching 40–50 Å, which indicated complete dissociation of the ligand from the binding site. Based on the RMSD analysis, hit compounds **SH2** and **SH5** were excluded and not considered for further investigations.

Next, the free energy landscapes (FELs) of the three prioritized ELMO1 hit compounds **SH1, SH3**, and **SH4** were generated (Figure 5) to investigate the conformational states of the complexes and assess their energy minima. The FEL of the SH1/ELMO1 complex (Figure 5A) revealed a single, broad energy minimum. The relatively wide basin along the protein RMSD axis indicates moderate backbone flexibility of ELMO1 observed in the RMSD trajectory. However, the unimodal character of the landscape confirms that these fluctuations occur within a single thermodynamic state rather than representing transitions between distinct conformational ensembles. Overall, the FEL of **SH1** is consistent with a stable binding mode.

**Figure 5.**
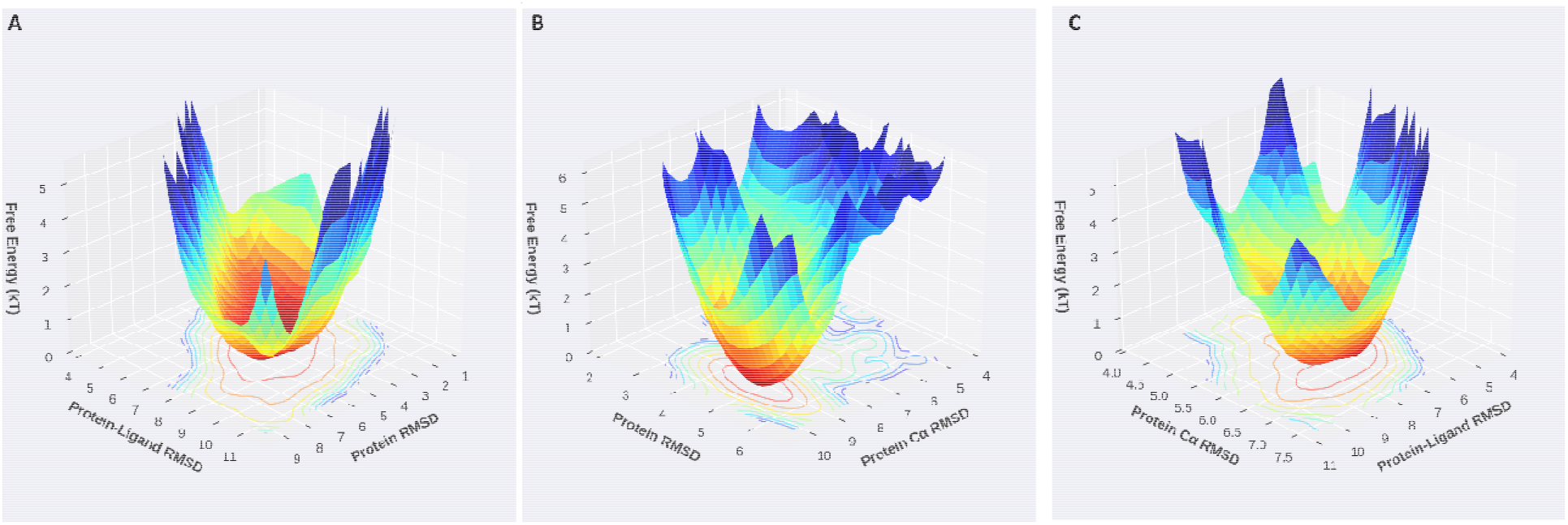
Free energy landscapes of prioritized ELMO1–hit compound complexes. Three-dimensional free energy landscapes (FELs) are shown for (A) **SH1**/ELMO1, (B) **SH3**/ELMO1, and (C) **SH4**/ELMO1 complexes.

The FEL of the **SH3**/ELMO1 complex (Figure 5B) showed a primary low-energy basin alongside several adjacent shallow sub-minima distributed across a broader range of protein RMSD values. The presence of secondary minima separated by low free energy barriers, indicates that the **SH3**/ELMO1 complex samples multiple distinct conformational substates during the simulation. Despite this conformational multiplicity, **SH3** displayed the lowest RMSD standard deviation of all five systems, which suggests that the transitions between sub-minima are subtle and do not involve large-scale structural rearrangements. As such, the FEL of **SH3** may be the reflection of a productive conformational sampling within a well-defined binding envelope rather than instability, which could be advantageous for induced-fit binding adaptability.

Lastly, the FEL of the **SH4**/ELMO1 complex (Figure 5C) exhibits the most well-defined and deeply funneled energy landscape among the three systems, with a single sharp global minimum and steep surrounding energy walls across both collective variable axes. The basin is narrow and centered at low values of both protein Cα RMSD and protein-ligand RMSD, indicating that the ELMO1/**SH4** complex is confined to a highly restricted region of conformational space throughout the simulation, which is a hallmark of thermodynamically stable ligand–protein interactions. Collectively, the FEL analysis indicates that all three predicted hits are predicted to maintain a stable complex with ELMO1.

Lastly, the binding free energies for the three prioritized ELMO1/hit complexes were estimated using the MM-GBSA (Figure 6). **SH3** displayed the most favorable binding free energies, where it frequently reached −55 to −65 kcal/mol, which indicates a strong predicted affinity for the ELMO1 binding pocket. However, the **SH3** MMGBSA profile exhibited the highest fluctuations, where it showed a periodic rise toward 0 kcal/mol, which likely corresponds to the transient conformational sub-states identified in its FEL. **SH1** demonstrated consistently negative MM-GBSA values throughout the entire trajectory, ranging from approximately −20 to −55 kcal/mol with a stable mean of approximately −35 to −40 kcal/mol and without the sharp fluctuations that were a characteristic of **SH3**. This sustained negative profile across 100 ns is indicative of a persistently engaged binding mode, which, when taken together with its unimodal FEL and acceptable RMSD convergence, reinforces **SH1** as a well-balanced hit with reliable predicted affinity. In contrast, **SH4**, which previously exhibited favorable RMSD and FEL profiles, exhibited MM-GBSA values clustering near 0 kcal/mol for much of the trajectory, where the negative values were observed during the initial 20 ns of simulation. Collectively, the MM-GBSA analysis results when combined with the RMSD and FEL analysis establish **SH1** and **SH3** as the primary candidates for future investigations and possible experimental validation.

**Figure 6.**
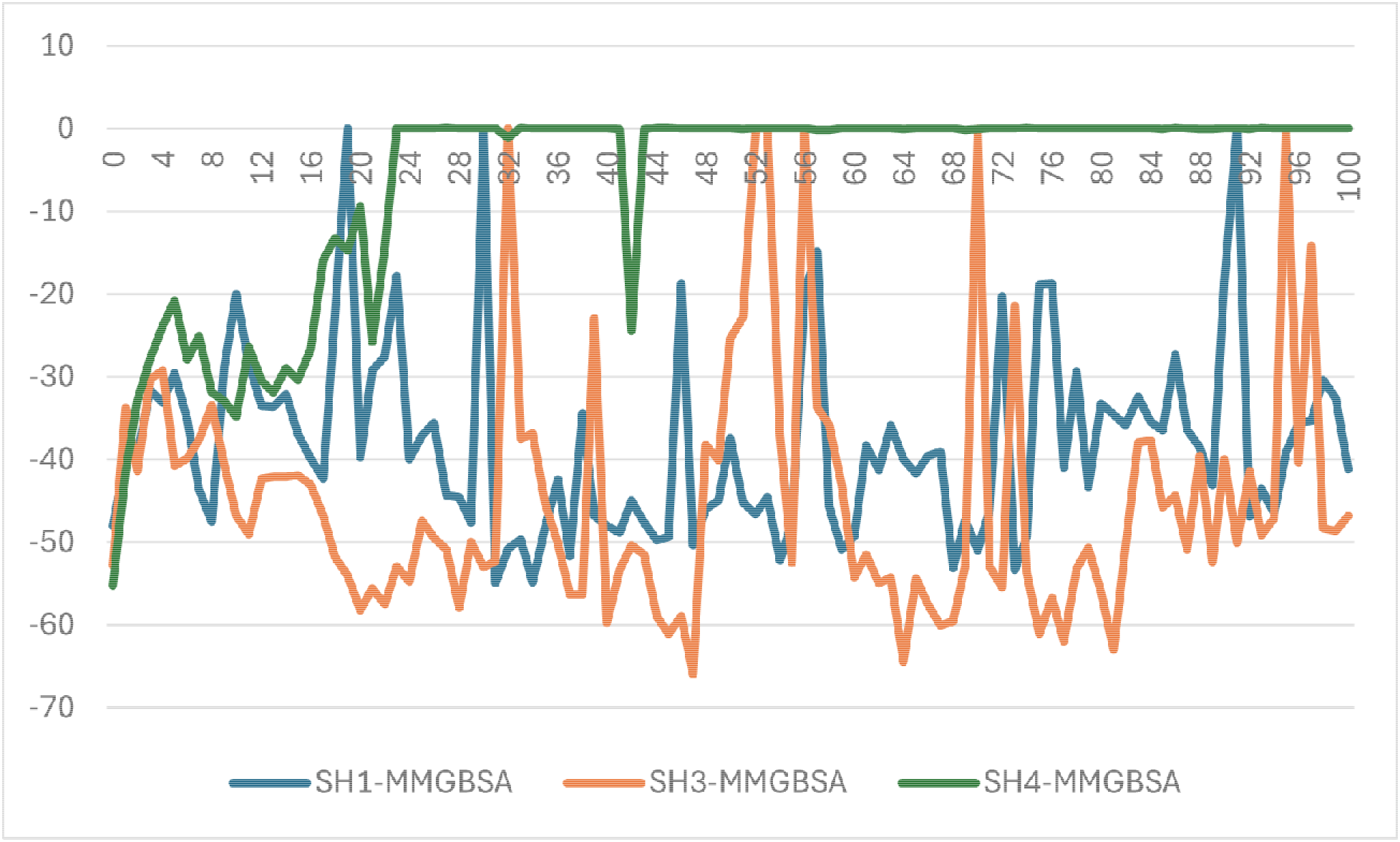
MM-GBSA binding free energies of hit compounds SH1, 3 and 5 4 in complex with ELMO1. Time-dependent MM-GBSA binding free energies (ΔG_bind, kcal/mol) are shown for **SH1**/ELMO1 (blue), **SH3**/ELMO1 (orange), and **SH4**/ELMO1 (green) complexes.

It should be noted that the results of our work are in silico in nature and should be subjected to experimental validation, where our lab is currently working on establishing the relevant models to do so. This work was designed to establish a foundation for future experimental studies targeting ELMO1 by acting as a starting point for hit-to-lead optimization campaigns, structure-activity relationship studies, and ultimately the development of first-in-class ELMO1 targeted inhibitors.

## 4. Conclusion

To the best of our knowledge, this study presents the first computational framework for the identification and characterization of small-molecule inhibitors targeting the ELMO1 binding interface aimed at disrupting the ELMO1/DOCK2 signaling axis, which has been implicated in various therapeutic conditions. Herein, we employed an in-house PPI.py script to analyze an MD simulation of the ELMO1/DOCK2 complex, which revealed a previously underexplored “bilobal” binding pocket on ELMO1 that represents a potential site for small molecules. From a focused library of 15,000 Enamine Diversity Discovery Set compounds, hierarchical virtual screening and MD-based post-docking validation predicted **SH1** and **SH3** as the most promising lead candidates, which were supported by favorable binding modes, sustained MM-GBSA binding free energies and thermodynamically stable free energy landscapes. Collectively, these findings lay a foundation for future experimental efforts toward the development of ELMO1-targeted inhibitors. Beyond the therapeutic potential of this work, we present two Python scripts: PPI.py and MD.py for analysis of Desmond-based MD of PPI and protein-ligand interactions, which have the potential to serve the whole drug discovery community.

## Supporting information

Supplemental raw data

## Notes

The authors declare no competing financial interests.

## Data sharing and code sharing

Raw MMHBSA data as well as the two python tools: MD.py and PPI.py can be found in the supplementary file.

