## Supplemental raw data for "A Computational Foundation Toward Targeting the ELMO1/DOCK2 Complex": MD.py

```
#!/usr/bin/env python3

import numpy as np
import pandas as pd
import matplotlib.pyplot as plt
import seaborn as sns
from scipy.stats import gaussian_kde
from scipy.ndimage import gaussian_filter, minimum_filter
import os
import sys
import warnings
warnings.filterwarnings('ignore')

# Set publication-quality style
plt.style.use('seaborn-v0_8-darkgrid')
sns.set_palette("husl")
plt.rcParams['figure.dpi'] = 100
plt.rcParams['savefig.dpi'] = 300
plt.rcParams['font.size'] = 11
plt.rcParams['axes.labelsize'] = 12
plt.rcParams['axes.titlesize'] = 14
plt.rcParams['xtick.labelsize'] = 10
plt.rcParams['ytick.labelsize'] = 10
plt.rcParams['legend.fontsize'] = 10

def read_data_file(filename):
    """Safely read Desmond output files with comprehensive error handling"""
    try:
        if not os.path.exists(filename):
            print(f"  ✗ File not found: {filename}")
            return None
            
        # Check file size
        if os.path.getsize(filename) == 0:
            print(f"  ✗ {filename} is empty")
            return None
        
        # Read first few lines to detect structure
        with open(filename, 'r') as f:
            first_lines = [f.readline() for _ in range(min(10, sum(1 for _ in f) + 10))]
        
        # Skip comment lines and find where data starts
        skip_rows = 0
        for i, line in enumerate(first_lines):
            if line.strip().startswith('#') or not line.strip():
                skip_rows = i + 1
            else:
                # Check if this line has numeric data
                try:
                    float(line.strip().split()[0])
                    skip_rows = i
                    break
                except (ValueError, IndexError):
                    skip_rows = i + 1
        
        # Read the file
        data = pd.read_csv(filename, sep=r'\s+', skiprows=skip_rows, header=None, 
                          engine='python', on_bad_lines='skip')
        
        # Check if we got actual data
        if data.empty or data.shape[0] == 0:
            print(f"  ✗ {filename} is empty or has no data")
            return None
            
        # Convert to numeric
        for col in data.columns:
            data[col] = pd.to_numeric(data[col], errors='coerce')
        
        # Remove rows with NaN
        data = data.dropna()
        
        if data.empty:
            print(f"  ✗ {filename} has no valid numeric data")
            return None
        
        print(f"  ✓ Loaded {filename}: {data.shape[0]} rows, {data.shape[1]} columns")
        return data
        
    except FileNotFoundError:
        print(f"  ✗ File not found: {filename}")
        return None
    except Exception as e:
        print(f"  ✗ Error reading {filename}: {str(e)}")
        return None

def plot_rmsd_analysis():
    """Comprehensive RMSD analysis with improved error handling"""
    print("\n[1] RMSD Analysis...")
    data = read_data_file('PL_RMSD.dat')
    if data is None or data.shape[1] < 2:
        print("  ⚠ Skipping RMSD analysis - insufficient data")
        return
    
    time = data.iloc[:, 0].values
    rmsd = data.iloc[:, 1].values
    
    # Validate data
    if len(rmsd) < 10:
        print("  ⚠ Too few data points for meaningful analysis")
        return
    
    fig = plt.figure(figsize=(14, 10))
    gs = fig.add_gridspec(3, 2, hspace=0.3, wspace=0.3)
    
    # 1. Time series
    ax1 = fig.add_subplot(gs[0, :])
    ax1.plot(time, rmsd, linewidth=1.5, color='#2E86AB', alpha=0.8)
    
    # Rolling average with dynamic window
    window = max(10, min(50, len(rmsd)//10))
    if window >= 3:
        rmsd_smooth = pd.Series(rmsd).rolling(window=window, center=True, min_periods=1).mean()
        ax1.plot(time, rmsd_smooth, linewidth=2.5, color='#E63946', 
                 label=f'Rolling avg (window={window})')
    
    mean_rmsd = np.mean(rmsd)
    std_rmsd = np.std(rmsd)
    ax1.axhline(y=mean_rmsd, color='green', linestyle='--', alpha=0.7, 
                label=f'Mean: {mean_rmsd:.2f} Å')
    ax1.fill_between(time, mean_rmsd-std_rmsd, mean_rmsd+std_rmsd, 
                      alpha=0.2, color='green', label=f'±1σ: {std_rmsd:.2f} Å')
    
    ax1.set_xlabel('Time (ns)', fontweight='bold')
    ax1.set_ylabel('RMSD (Å)', fontweight='bold')
    ax1.set_title('Protein-Ligand Complex RMSD Over Time', fontweight='bold', pad=10)
    ax1.legend(loc='best')
    ax1.grid(alpha=0.3)
    
    # 2. Distribution
    ax2 = fig.add_subplot(gs[1, 0])
    n_bins = min(40, max(10, len(rmsd)//25))
    counts, bins, patches = ax2.hist(rmsd, bins=n_bins, density=True, alpha=0.7, 
                                     color='#3498DB', edgecolor='black')
    
    # KDE overlay
    try:
        kde = gaussian_kde(rmsd)
        x_range = np.linspace(rmsd.min(), rmsd.max(), 200)
        ax2.plot(x_range, kde(x_range), 'r-', linewidth=2.5, label='KDE')
    except:
        pass
    
    ax2.axvline(mean_rmsd, color='green', linestyle='--', linewidth=2, label='Mean')
    
    ax2.set_xlabel('RMSD (Å)', fontweight='bold')
    ax2.set_ylabel('Probability Density', fontweight='bold')
    ax2.set_title('RMSD Distribution', fontweight='bold')
    ax2.legend()
    ax2.grid(alpha=0.3)
    
    # 3. Free energy profile (1D)
    ax3 = fig.add_subplot(gs[1, 1])
    fe_bins = min(50, max(20, len(rmsd)//20))
    counts, bins = np.histogram(rmsd, bins=fe_bins, density=True)
    bin_centers = (bins[:-1] + bins[1:]) / 2
    
    # Avoid log(0)
    counts_safe = np.where(counts > 0, counts, np.nan)
    free_energy = -np.log(counts_safe)
    free_energy = free_energy - np.nanmin(free_energy)
    
    # Filter out NaN values
    valid_mask = ~np.isnan(free_energy)
    ax3.plot(bin_centers[valid_mask], free_energy[valid_mask], 'o-', linewidth=2, markersize=6, 
             color='#E74C3C', markerfacecolor='white', markeredgewidth=2)
    ax3.fill_between(bin_centers[valid_mask], free_energy[valid_mask], 
                      alpha=0.3, color='#E74C3C')
    
    ax3.set_xlabel('RMSD (Å)', fontweight='bold')
    ax3.set_ylabel('Free Energy (kT)', fontweight='bold')
    ax3.set_title('1D Free Energy Profile', fontweight='bold')
    ax3.grid(alpha=0.3)
    
    # 4. Cumulative distribution
    ax4 = fig.add_subplot(gs[2, 0])
    sorted_rmsd = np.sort(rmsd)
    cumulative = np.arange(1, len(sorted_rmsd) + 1) / len(sorted_rmsd)
    ax4.plot(sorted_rmsd, cumulative, linewidth=2.5, color='#9B59B6')
    ax4.axhline(0.5, color='red', linestyle='--', alpha=0.7)
    ax4.axvline(np.median(rmsd), color='red', linestyle='--', alpha=0.7,
                label=f'Median: {np.median(rmsd):.2f} Å')
    
    ax4.set_xlabel('RMSD (Å)', fontweight='bold')
    ax4.set_ylabel('Cumulative Probability', fontweight='bold')
    ax4.set_title('Cumulative Distribution Function', fontweight='bold')
    ax4.legend()
    ax4.grid(alpha=0.3)
    
    # 5. Statistics box
    ax5 = fig.add_subplot(gs[2, 1])
    ax5.axis('off')
    
    stats_text = f"""
    RMSD Statistics
    ───────────────────────
    Mean:           {mean_rmsd:.3f} Å
    Median:         {np.median(rmsd):.3f} Å
    Std Dev:        {std_rmsd:.3f} Å
    Min:            {np.min(rmsd):.3f} Å
    Max:            {np.max(rmsd):.3f} Å
    
    Quartiles:
    Q1 (25%):       {np.percentile(rmsd, 25):.3f} Å
    Q2 (50%):       {np.percentile(rmsd, 50):.3f} Å
    Q3 (75%):       {np.percentile(rmsd, 75):.3f} Å
    
    Simulation Time: {time[-1]:.2f} ns
    Total Frames:    {len(rmsd)}
    """
    
    ax5.text(0.1, 0.9, stats_text, transform=ax5.transAxes,
             fontsize=10, verticalalignment='top', fontfamily='monospace',
             bbox=dict(boxstyle='round', facecolor='wheat', alpha=0.5))
    
    plt.savefig('RMSD_comprehensive_analysis.png', dpi=300, bbox_inches='tight')
    plt.close()
    print("  ✓ Saved: RMSD_comprehensive_analysis.png")

def create_2d_fel():
    """Create publication-quality 2D and 3D Free Energy Landscapes"""
    print("\n[2] Creating Free Energy Landscapes...")
    
    rmsd_data = read_data_file('PL_RMSD.dat')
    if rmsd_data is None or rmsd_data.shape[1] < 2:
        print("  ⚠ Skipping FEL - insufficient RMSD data")
        return
    
    # Extract available columns
    time = rmsd_data.iloc[:, 0].values
    
    print(f"  → RMSD file has {rmsd_data.shape[1]} columns")
    print(f"  → Column ranges: ", end="")
    for i in range(min(5, rmsd_data.shape[1])):
        col_data = rmsd_data.iloc[:, i].values
        print(f"Col{i}[{np.min(col_data):.2f}-{np.max(col_data):.2f}] ", end="")
    print()
    
    # Determine which columns to use for FEL - be smarter about it
    # Column 0 is always time, skip it
    # Look for columns with RMSD-like values (typically 0.5 to 10 Å)
    
    valid_rmsd_cols = []
    for i in range(1, rmsd_data.shape[1]):
        col_data = rmsd_data.iloc[:, i].values
        col_min, col_max = np.min(col_data), np.max(col_data)
        col_mean = np.mean(col_data)
        col_std = np.std(col_data)
        
        # RMSD values are typically between 0 and 15 Å, mean usually 1-5 Å
        if 0 <= col_min < 20 and col_max < 20 and 0.1 < col_mean < 15:
            valid_rmsd_cols.append((i, col_mean, col_std, col_max - col_min))
            print(f"  → Column {i} looks like RMSD: mean={col_mean:.2f}±{col_std:.2f} Å, range=[{col_min:.2f}, {col_max:.2f}]")
    
    if len(valid_rmsd_cols) >= 2:
        # Sort by standard deviation (variance) to get most different columns
        # We want columns that show different dynamics
        valid_rmsd_cols_sorted = sorted(valid_rmsd_cols, key=lambda x: x[2], reverse=True)
        
        # Pick two columns with most different characteristics
        # Strategy: use the column with highest variance and one that's most different
        col1_idx = valid_rmsd_cols_sorted[0][0]  # Highest variance
        
        # Find column most different from col1
        col1_data = rmsd_data.iloc[:, col1_idx].values
        max_diff = 0
        col2_idx = valid_rmsd_cols_sorted[1][0] if len(valid_rmsd_cols_sorted) > 1 else col1_idx
        
        for idx, mean, std, rng in valid_rmsd_cols_sorted[1:]:
            col_data = rmsd_data.iloc[:, idx].values
            # Calculate correlation - lower is better (more different)
            correlation = np.corrcoef(col1_data, col_data)[0, 1]
            diff_score = (1 - abs(correlation)) * std  # Prefer low correlation and high variance
            
            if diff_score > max_diff:
                max_diff = diff_score
                col2_idx = idx
        
        rmsd_col1 = rmsd_data.iloc[:, col1_idx].values
        rmsd_col2 = rmsd_data.iloc[:, col2_idx].values
        
        correlation = np.corrcoef(rmsd_col1, rmsd_col2)[0, 1]
        print(f"  → Selected columns {col1_idx} vs {col2_idx} (correlation: {correlation:.3f})")
        
        # Try to intelligently label based on typical Desmond output
        # Columns usually: 1=Protein-Ligand, 2=Ligand, 3=Protein Cα, 4=Protein backbone
        labels_map = {
            1: 'Protein-Ligand RMSD',
            2: 'Ligand RMSD', 
            3: 'Protein Cα RMSD',
            4: 'Protein Backbone RMSD',
            5: 'Protein RMSD',
            6: 'Protein Side-chain RMSD'
        }
        
        coord1_label = f'{labels_map.get(col1_idx, f"RMSD Col{col1_idx}")} (Å)'
        coord2_label = f'{labels_map.get(col2_idx, f"RMSD Col{col2_idx}")} (Å)'
        
    elif len(valid_rmsd_cols) == 1:
        # Only one valid RMSD column - use it vs its derivative
        col_idx = valid_rmsd_cols[0][0]
        rmsd_col1 = rmsd_data.iloc[:, col_idx].values
        rmsd_col2 = np.gradient(rmsd_col1)
        coord1_label = 'RMSD (Å)'
        coord2_label = 'RMSD Change Rate (Å/frame)'
        print(f"  → Using column {col_idx} and its derivative")
        
    else:
        # Fallback - just use columns 1 and 2 or 1 and derivative
        print(f"  ⚠ Warning: Could not identify RMSD columns clearly")
        if rmsd_data.shape[1] >= 3:
            rmsd_col1 = rmsd_data.iloc[:, 1].values
            rmsd_col2 = rmsd_data.iloc[:, 2].values
            coord1_label = 'Column 1 (Å)'
            coord2_label = 'Column 2 (Å)'
        else:
            rmsd_col1 = rmsd_data.iloc[:, 1].values
            rmsd_col2 = np.gradient(rmsd_col1)
            coord1_label = 'RMSD (Å)'
            coord2_label = 'RMSD Change Rate (Å/frame)'
    
    rmsd = rmsd_col1  # For compatibility with contact analysis
    
    # Additional validation - check for reasonable ranges
    if np.max(rmsd_col1) > 50 or np.max(np.abs(rmsd_col2)) > 50:
        print(f"  ⚠ Warning: Unusual data ranges detected!")
        print(f"     Col1 range: [{np.min(rmsd_col1):.2f}, {np.max(rmsd_col1):.2f}]")
        print(f"     Col2 range: [{np.min(rmsd_col2):.2f}, {np.max(rmsd_col2):.2f}]")
    
    # Validate data
    if len(rmsd_col1) < 50:
        print("  ⚠ Too few data points for FEL - need at least 50 frames")
        return
    
    # Higher resolution for smoother FEL
    bins = min(100, max(30, int(np.sqrt(len(rmsd_col1)))))
    
    try:
        # Remove outliers for better binning (use 1st-99th percentile range)
        p1_x, p99_x = np.percentile(rmsd_col1, [1, 99])
        p1_y, p99_y = np.percentile(rmsd_col2, [1, 99])
        
        # Create bins focused on the actual data range
        x_range = [p1_x - 0.1*(p99_x-p1_x), p99_x + 0.1*(p99_x-p1_x)]
        y_range = [p1_y - 0.1*(p99_y-p1_y), p99_y + 0.1*(p99_y-p1_y)]
        
        print(f"  → Data range X: [{x_range[0]:.2f}, {x_range[1]:.2f}] Å")
        print(f"  → Data range Y: [{y_range[0]:.2f}, {y_range[1]:.2f}] Å")
        
        # Calculate 2D histogram using actual data columns with focused range
        H, xedges, yedges = np.histogram2d(rmsd_col1, rmsd_col2, bins=bins, 
                                           range=[x_range, y_range])
        
        # Apply Gaussian smoothing
        H_smooth = gaussian_filter(H, sigma=1.5)
        
        # Mask very low probability regions (less than 0.1% of max)
        threshold = 0.001 * np.max(H_smooth)
        H_smooth = np.ma.masked_where(H_smooth < threshold, H_smooth)
        
        # Calculate free energy with proper normalization
        # Convert counts to probability
        H_prob = H_smooth / np.sum(H_smooth)
        
        # Calculate free energy: ΔG = -RT ln(P/P_ref) = -kT ln(P/P_max)
        # Setting P_ref = P_max makes the minimum energy = 0
        free_energy = -np.log(H_prob / np.ma.max(H_prob))
        
        # Convert from natural log units to kT (already in kT since we used ln)
        # Cap at reasonable maximum
        vmax = min(np.percentile(free_energy.compressed(), 95), 8)
        
        print(f"  → Free energy range: [0.00, {vmax:.2f}] kT")
        
        # Create mesh grids
        X = (xedges[:-1] + xedges[1:]) / 2
        Y = (yedges[:-1] + yedges[1:]) / 2
        X_mesh, Y_mesh = np.meshgrid(X, Y)
        
        # Create figure with better spacing
        fig = plt.figure(figsize=(20, 6.5))
        
        # ========== 2D Contour Plot with Energy Basins ==========
        ax1 = plt.subplot(1, 3, 1)
        
        # Use inverted colormap (blue=low energy, red=high energy)
        levels_filled = np.linspace(0, vmax, 30)
        im1 = ax1.contourf(X_mesh, Y_mesh, free_energy.T, levels=levels_filled,
                           cmap='jet_r', extend='max', antialiased=True)
        
        # Add bold contour lines to distinguish basins
        contour_levels = [0.5, 1.0, 1.5, 2.0, 2.5, 3.0, 4.0, 5.0, 6.0]
        contour_levels = [c for c in contour_levels if c < vmax]
        if len(contour_levels) > 0:
            contours = ax1.contour(X_mesh, Y_mesh, free_energy.T, levels=contour_levels,
                                  colors='black', linewidths=1.5, alpha=0.7)
            ax1.clabel(contours, inline=True, fontsize=10, fmt='%1.1f')
        
        # Find and mark local minima (but only in low energy regions < 2 kT)
        min_filtered = minimum_filter(free_energy.filled(999), size=10)
        local_minima = (free_energy == min_filtered) & (free_energy < 2.0)
        
        min_coords = np.where(local_minima)
        minima_found = []
        
        if len(min_coords[0]) > 0:
            # Sort by energy value
            for i in range(len(min_coords[0])):
                ix, iy = min_coords[0][i], min_coords[1][i]
                energy_val = free_energy[ix, iy]
                minima_found.append((ix, iy, energy_val))
            
            minima_found.sort(key=lambda x: x[2])  # Sort by energy
            
            # Plot top 3 minima
            for i, (ix, iy, energy_val) in enumerate(minima_found[:3]):
                if i == 0:
                    ax1.plot(X[ix], Y[iy], 'w*', markersize=28, markeredgecolor='black',
                            markeredgewidth=2.5, label=f'Global min: {energy_val:.2f} kT', zorder=10)
                elif i == 1:
                    ax1.plot(X[ix], Y[iy], 'y*', markersize=22, markeredgecolor='black',
                            markeredgewidth=2, label=f'Local min: {energy_val:.2f} kT', zorder=10)
                else:
                    ax1.plot(X[ix], Y[iy], 'y*', markersize=18, markeredgecolor='black',
                            markeredgewidth=1.5, zorder=10)
        
        cbar1 = plt.colorbar(im1, ax=ax1, pad=0.02, fraction=0.046)
        cbar1.set_label('Free Energy (kT)', fontsize=12, fontweight='bold')
        
        ax1.set_xlabel(coord1_label, fontsize=12, fontweight='bold')
        ax1.set_ylabel(coord2_label, fontsize=12, fontweight='bold')
        ax1.set_title('2D Free Energy Landscape', fontsize=13, fontweight='bold', pad=10)
        ax1.legend(loc='best', fontsize=9)
        ax1.grid(alpha=0.2)
        ax1.set_aspect('auto')
        
        # ========== 3D Surface Plot ==========
        ax2 = plt.subplot(1, 3, 2, projection='3d')
        
        # Plot 3D surface
        surf = ax2.plot_surface(X_mesh, Y_mesh, free_energy.T, cmap='jet_r',
                               vmin=0, vmax=vmax, alpha=0.9, 
                               edgecolor='none', antialiased=True, shade=True)
        
        # Add contour projections on bottom
        ax2.contour(X_mesh, Y_mesh, free_energy.T, levels=10, 
                   zdir='z', offset=0, cmap='jet_r', linewidths=1, alpha=0.5)
        
        # Split long labels for 3D plot
        xlabel_3d = coord1_label.split('(')[0].strip()
        ylabel_3d = coord2_label.split('(')[0].strip()
        
        ax2.set_xlabel(xlabel_3d, fontsize=11, fontweight='bold', labelpad=8)
        ax2.set_ylabel(ylabel_3d, fontsize=11, fontweight='bold', labelpad=8)
        ax2.set_zlabel('Free Energy (kT)', fontsize=11, fontweight='bold', labelpad=8)
        ax2.set_title('3D Free Energy Surface', fontsize=13, fontweight='bold', pad=10)
        ax2.view_init(elev=25, azim=45)
        ax2.set_zlim(0, vmax)
        ax2.set_box_aspect(None, zoom=0.85)
        
        # ========== FEL with Contacts (if available) ==========
        ax3 = plt.subplot(1, 3, 3)
        
        contact_file = None
        for f in ['PL-Contacts_HBond.dat', 'PL-Contacts_Hydrophobic.dat', 
                  'PL-Contacts_Ionic.dat', 'PL-Contacts_WaterBridge.dat']:
            if os.path.exists(f):
                contact_file = f
                break
        
        if contact_file:
            contact_data = read_data_file(contact_file)
            if contact_data is not None and contact_data.shape[1] >= 2:
                contacts = contact_data.iloc[:, 1].values
                min_len = min(len(rmsd), len(contacts))
                
                if min_len >= 50:
                    # Use same focused range approach
                    p1_x2, p99_x2 = np.percentile(rmsd[:min_len], [1, 99])
                    p1_y2, p99_y2 = np.percentile(contacts[:min_len], [1, 99])
                    
                    x_range2 = [p1_x2 - 0.1*(p99_x2-p1_x2), p99_x2 + 0.1*(p99_x2-p1_x2)]
                    y_range2 = [max(0, p1_y2 - 0.5), p99_y2 + 0.5]  # Contacts should be >= 0
                    
                    H2, xedges2, yedges2 = np.histogram2d(rmsd[:min_len], contacts[:min_len], 
                                                          bins=bins, range=[x_range2, y_range2])
                    H2_smooth = gaussian_filter(H2, sigma=1.5)
                    
                    threshold2 = 0.001 * np.max(H2_smooth)
                    H2_smooth = np.ma.masked_where(H2_smooth < threshold2, H2_smooth)
                    
                    # Proper free energy calculation
                    H2_prob = H2_smooth / np.sum(H2_smooth)
                    free_energy2 = -np.log(H2_prob / np.ma.max(H2_prob))
                    vmax2 = min(np.percentile(free_energy2.compressed(), 95), 8)
                    
                    X2 = (xedges2[:-1] + xedges2[1:]) / 2
                    Y2 = (yedges2[:-1] + yedges2[1:]) / 2
                    X2_mesh, Y2_mesh = np.meshgrid(X2, Y2)
                    
                    im3 = ax3.contourf(X2_mesh, Y2_mesh, free_energy2.T, levels=30,
                                      cmap='jet_r', extend='max', antialiased=True)
                    
                    contour_levels3 = [c for c in [0.5, 1.0, 1.5, 2.0, 2.5, 3.0, 4.0, 5.0] if c < vmax2]
                    if len(contour_levels3) > 0:
                        contours3 = ax3.contour(X2_mesh, Y2_mesh, free_energy2.T,
                                               levels=contour_levels3,
                                               colors='black', linewidths=1.5, alpha=0.7)
                        ax3.clabel(contours3, inline=True, fontsize=10, fmt='%1.1f')
                    
                    # Find minima
                    min_filtered2 = minimum_filter(free_energy2.filled(999), size=10)
                    local_minima2 = (free_energy2 == min_filtered2) & (free_energy2 < 2.0)
                    min_coords2 = np.where(local_minima2)
                    
                    if len(min_coords2[0]) > 0:
                        ix2, iy2 = min_coords2[0][0], min_coords2[1][0]
                        ax3.plot(X2[ix2], Y2[iy2], 'w*', markersize=25, 
                                markeredgecolor='black', markeredgewidth=2,
                                label=f'Global min: {free_energy2[ix2, iy2]:.2f} kT')
                    
                    cbar3 = plt.colorbar(im3, ax=ax3, pad=0.02, fraction=0.046)
                    cbar3.set_label('Free Energy (kT)', fontsize=12, fontweight='bold')
                    
                    contact_type = contact_file.replace('PL-Contacts_', '').replace('.dat', '')
                    ax3.set_xlabel(coord1_label.split()[0] + ' (Å)', fontsize=12, fontweight='bold')
                    ax3.set_ylabel(f'{contact_type} Contacts', fontsize=12, fontweight='bold')
                    ax3.set_title(f'FEL: RMSD vs {contact_type}', fontsize=13, fontweight='bold', pad=10)
                    ax3.legend(loc='best', fontsize=9)
                    ax3.grid(alpha=0.2)
                    ax3.set_aspect('auto')
                else:
                    ax3.text(0.5, 0.5, 'Insufficient contact data',
                            ha='center', va='center', fontsize=14,
                            bbox=dict(boxstyle='round', facecolor='wheat', alpha=0.7))
                    ax3.axis('off')
            else:
                ax3.text(0.5, 0.5, 'Contact data not available',
                        ha='center', va='center', fontsize=14,
                        bbox=dict(boxstyle='round', facecolor='wheat', alpha=0.7))
                ax3.axis('off')
        else:
            ax3.text(0.5, 0.5, 'No contact data found\n\nLooking for:\n• PL-Contacts_HBond.dat\n• PL-Contacts_Hydrophobic.dat\n• PL-Contacts_Ionic.dat',
                    ha='center', va='center', fontsize=11,
                    bbox=dict(boxstyle='round', facecolor='lightcoral', alpha=0.5))
            ax3.axis('off')
        
        plt.tight_layout()
        plt.savefig('FEL_2D_landscapes.png', dpi=300, bbox_inches='tight')
        plt.close()
        print("  ✓ Saved: FEL_2D_landscapes.png")
        
    except Exception as e:
        print(f"  ✗ Error creating FEL: {str(e)}")
        import traceback
        traceback.print_exc()

def plot_contact_analysis():
    """Comprehensive contact analysis with improved error handling"""
    print("\n[3] Contact Analysis...")
    
    contact_files = {
        'H-Bonds': ('PL-Contacts_HBond.dat', '#E63946'),
        'Hydrophobic': ('PL-Contacts_Hydrophobic.dat', '#F1C40F'),
        'Ionic': ('PL-Contacts_Ionic.dat', '#2ECC71'),
        'Water Bridges': ('PL-Contacts_WaterBridge.dat', '#3498DB'),
        'Pi-Pi': ('PL-Contacts_Pi-Pi.dat', '#9B59B6'),
        'Pi-Cation': ('PL-Contacts_Pi-Cation.dat', '#E67E22')
    }
    
    contact_data_dict = {}
    
    # Load all contact data
    for label, (file, color) in contact_files.items():
        data = read_data_file(file)
        if data is not None and data.shape[1] >= 2:
            contact_data_dict[label] = {
                'time': data.iloc[:, 0].values,
                'contacts': data.iloc[:, 1].values,
                'color': color
            }
    
    if not contact_data_dict:
        print("  ⚠ No contact data available - skipping contact analysis")
        return
    
    fig, axes = plt.subplots(2, 2, figsize=(15, 10))
    axes = axes.flatten()
    
    # Plot 1: All contacts over time
    ax1 = axes[0]
    for label, data_dict in contact_data_dict.items():
        ax1.plot(data_dict['time'], data_dict['contacts'], 
                label=label, linewidth=2, color=data_dict['color'], alpha=0.8)
    
    ax1.set_xlabel('Time (ns)', fontweight='bold')
    ax1.set_ylabel('Number of Contacts', fontweight='bold')
    ax1.set_title('Protein-Ligand Interactions Over Time', fontweight='bold', pad=10)
    ax1.legend(loc='best', frameon=True, shadow=True)
    ax1.grid(alpha=0.3)
    
    # Plot 2: Average contacts bar chart
    ax2 = axes[1]
    labels = list(contact_data_dict.keys())
    means = [np.mean(contact_data_dict[label]['contacts']) for label in labels]
    stds = [np.std(contact_data_dict[label]['contacts']) for label in labels]
    colors = [contact_data_dict[label]['color'] for label in labels]
    
    bars = ax2.bar(range(len(labels)), means, yerr=stds, 
                  color=colors, alpha=0.7, capsize=5, edgecolor='black', linewidth=1.5)
    ax2.set_xticks(range(len(labels)))
    ax2.set_xticklabels(labels, rotation=45, ha='right')
    ax2.set_ylabel('Average Number of Contacts', fontweight='bold')
    ax2.set_title('Average Contact Counts', fontweight='bold', pad=10)
    ax2.grid(axis='y', alpha=0.3)
    
    # Add value labels on bars
    for i, (bar, mean) in enumerate(zip(bars, means)):
        height = bar.get_height()
        ax2.text(bar.get_x() + bar.get_width()/2., height,
                f'{mean:.1f}', ha='center', va='bottom', fontsize=9)
    
    # Plot 3: Contact persistence
    ax3 = axes[2]
    for i, label in enumerate(labels):
        data = contact_data_dict[label]['contacts']
        unique, counts = np.unique(data.astype(int), return_counts=True)
        persistence = counts / len(data) * 100
        ax3.bar(unique + i*0.1, persistence, width=0.1, 
               label=label, color=colors[i], alpha=0.7)
    
    ax3.set_xlabel('Number of Contacts', fontweight='bold')
    ax3.set_ylabel('Occurrence (%)', fontweight='bold')
    ax3.set_title('Contact Number Distribution', fontweight='bold', pad=10)
    ax3.legend(fontsize=8, ncol=2)
    ax3.grid(axis='y', alpha=0.3)
    
    # Plot 4: Statistics table
    ax4 = axes[3]
    ax4.axis('off')
    
    stats_lines = ["Contact Statistics\n" + "─"*30]
    for label in labels:
        contacts = contact_data_dict[label]['contacts']
        stats_lines.append(f"{label:15s} {np.mean(contacts):5.2f} ± {np.std(contacts):4.2f}")
    
    stats_text = '\n'.join(stats_lines)
    ax4.text(0.1, 0.9, stats_text, transform=ax4.transAxes,
             fontsize=10, verticalalignment='top', fontfamily='monospace',
             bbox=dict(boxstyle='round', facecolor='lightblue', alpha=0.5))
    
    plt.tight_layout()
    plt.savefig('Contact_comprehensive_analysis.png', dpi=300, bbox_inches='tight')
    plt.close()
    print("  ✓ Saved: Contact_comprehensive_analysis.png")

def plot_rmsf_analysis():
    """Enhanced RMSF visualization with improved error handling"""
    print("\n[4] RMSF Analysis...")
    
    p_rmsf = read_data_file('P_RMSF.dat')
    l_rmsf = read_data_file('L_RMSF.dat')
    
    if p_rmsf is None and l_rmsf is None:
        print("  ⚠ No RMSF data available - skipping")
        return
    
    n_plots = sum([p_rmsf is not None, l_rmsf is not None])
    fig, axes = plt.subplots(n_plots, 1, figsize=(14, 5*n_plots))
    
    if n_plots == 1:
        axes = [axes]
    
    plot_idx = 0
    
    # Protein RMSF
    if p_rmsf is not None and p_rmsf.shape[1] >= 2:
        residues = p_rmsf.iloc[:, 0].values
        rmsf = p_rmsf.iloc[:, 1].values
        
        # Create gradient color based on RMSF value
        colors = plt.cm.RdYlBu_r(rmsf / np.max(rmsf))
        
        axes[plot_idx].bar(residues, rmsf, color=colors, alpha=0.8, width=1.0, 
                          edgecolor='black', linewidth=0.3)
        
        # Add threshold line for flexible regions
        threshold = np.mean(rmsf) + np.std(rmsf)
        axes[plot_idx].axhline(y=threshold, color='red', linestyle='--', 
                              linewidth=2, label=f'Flexible threshold: {threshold:.2f} Å')
        axes[plot_idx].axhline(y=np.mean(rmsf), color='green', linestyle='--',
                              linewidth=2, label=f'Mean: {np.mean(rmsf):.2f} Å')
        
        axes[plot_idx].set_xlabel('Residue Number', fontsize=12, fontweight='bold')
        axes[plot_idx].set_ylabel('RMSF (Å)', fontsize=12, fontweight='bold')
        axes[plot_idx].set_title('Protein Residue Flexibility (RMSF)', 
                                fontsize=13, fontweight='bold', pad=10)
        axes[plot_idx].legend()
        axes[plot_idx].grid(alpha=0.3, axis='y')
        plot_idx += 1
    
    # Ligand RMSF
    if l_rmsf is not None and l_rmsf.shape[1] >= 2:
        atoms = l_rmsf.iloc[:, 0].values
        rmsf = l_rmsf.iloc[:, 1].values
        
        colors = plt.cm.plasma(rmsf / np.max(rmsf))
        
        axes[plot_idx].bar(atoms, rmsf, color=colors, alpha=0.8, width=0.7, 
                          edgecolor='black', linewidth=0.5)
        axes[plot_idx].axhline(y=np.mean(rmsf), color='red', linestyle='--',
                              linewidth=2, label=f'Mean: {np.mean(rmsf):.2f} Å')
        
        axes[plot_idx].set_xlabel('Atom Number', fontsize=12, fontweight='bold')
        axes[plot_idx].set_ylabel('RMSF (Å)', fontsize=12, fontweight='bold')
        axes[plot_idx].set_title('Ligand Atom Flexibility (RMSF)', 
                                fontsize=13, fontweight='bold', pad=10)
        axes[plot_idx].legend()
        axes[plot_idx].grid(alpha=0.3, axis='y')
    
    plt.tight_layout()
    plt.savefig('RMSF_enhanced_analysis.png', dpi=300, bbox_inches='tight')
    plt.close()
    print("  ✓ Saved: RMSF_enhanced_analysis.png")

def create_summary_report():
    """Create a comprehensive text summary report"""
    print("\n[5] Generating Summary Report...")
    
    report = []
    report.append("="*70)
    report.append("DESMOND MD SIMULATION ANALYSIS SUMMARY")
    report.append("="*70)
    report.append("")
    
    # RMSD stats
    rmsd_data = read_data_file('PL_RMSD.dat')
    if rmsd_data is not None and rmsd_data.shape[1] >= 2:
        rmsd = rmsd_data.iloc[:, 1].values
        time = rmsd_data.iloc[:, 0].values
        
        report.append("RMSD STATISTICS:")
        report.append("-" * 70)
        report.append(f"  Simulation Time:      {time[-1]:.2f} ns")
        report.append(f"  Total Frames:         {len(rmsd)}")
        report.append(f"  Mean RMSD:            {np.mean(rmsd):.3f} ± {np.std(rmsd):.3f} Å")
        report.append(f"  Median RMSD:          {np.median(rmsd):.3f} Å")
        report.append(f"  Min RMSD:             {np.min(rmsd):.3f} Å")
        report.append(f"  Max RMSD:             {np.max(rmsd):.3f} Å")
        report.append(f"  RMSD Range:           {np.max(rmsd) - np.min(rmsd):.3f} Å")
        
        # Equilibration check
        first_half = np.mean(rmsd[:len(rmsd)//2])
        second_half = np.mean(rmsd[len(rmsd)//2:])
        report.append(f"  First Half Mean:      {first_half:.3f} Å")
        report.append(f"  Second Half Mean:     {second_half:.3f} Å")
        report.append(f"  Difference:           {abs(second_half - first_half):.3f} Å")
        
        if abs(second_half - first_half) < np.std(rmsd):
            report.append("  ✓ System appears EQUILIBRATED")
        else:
            report.append("  ⚠ System may still be EQUILIBRATING")
        report.append("")
    
    # Contact stats
    report.append("INTERACTION STATISTICS:")
    report.append("-" * 70)
    
    contact_files = {
        'H-Bonds': 'PL-Contacts_HBond.dat',
        'Hydrophobic': 'PL-Contacts_Hydrophobic.dat',
        'Ionic': 'PL-Contacts_Ionic.dat',
        'Water Bridges': 'PL-Contacts_WaterBridge.dat',
        'Pi-Pi': 'PL-Contacts_Pi-Pi.dat',
        'Pi-Cation': 'PL-Contacts_Pi-Cation.dat'
    }
    
    for label, file in contact_files.items():
        data = read_data_file(file)
        if data is not None and data.shape[1] >= 2:
            contacts = data.iloc[:, 1].values
            report.append(f"  {label:20s} {np.mean(contacts):.2f} ± {np.std(contacts):.2f}")
    
    report.append("")
    
    # RMSF stats
    p_rmsf = read_data_file('P_RMSF.dat')
    if p_rmsf is not None and p_rmsf.shape[1] >= 2:
        rmsf = p_rmsf.iloc[:, 1].values
        report.append("PROTEIN FLEXIBILITY (RMSF):")
        report.append("-" * 70)
        report.append(f"  Mean RMSF:            {np.mean(rmsf):.3f} Å")
        report.append(f"  Max RMSF:             {np.max(rmsf):.3f} Å (Residue {int(p_rmsf.iloc[np.argmax(rmsf), 0])})")
        report.append(f"  Min RMSF:             {np.min(rmsf):.3f} Å (Residue {int(p_rmsf.iloc[np.argmin(rmsf), 0])})")
        
        # Identify flexible regions (> mean + 1 std)
        threshold = np.mean(rmsf) + np.std(rmsf)
        flexible_residues = p_rmsf.iloc[rmsf > threshold, 0].values.astype(int)
        if len(flexible_residues) > 0:
            report.append(f"  Highly Flexible:      {len(flexible_residues)} residues (>{threshold:.2f} Å)")
            if len(flexible_residues) <= 20:
                report.append(f"  Flexible Residues:    {', '.join(map(str, flexible_residues))}")
        report.append("")
    
    report.append("="*70)
    report.append("Generated files:")
    if os.path.exists('RMSD_comprehensive_analysis.png'):
        report.append("  ✓ RMSD_comprehensive_analysis.png")
    if os.path.exists('FEL_2D_landscapes.png'):
        report.append("  ✓ FEL_2D_landscapes.png")
    if os.path.exists('Contact_comprehensive_analysis.png'):
        report.append("  ✓ Contact_comprehensive_analysis.png")
    if os.path.exists('RMSF_enhanced_analysis.png'):
        report.append("  ✓ RMSF_enhanced_analysis.png")
    report.append("="*70)
    
    # Print to console
    for line in report:
        print(line)
    
    # Save to file
    with open('MD_Analysis_Summary.txt', 'w') as f:
        f.write('\n'.join(report))
    
    print("\n  ✓ Saved: MD_Analysis_Summary.txt")

# Main execution
if __name__ == "__main__":
    print("\n" + "="*70)
    print("COMPREHENSIVE DESMOND MD ANALYSIS - FINAL IMPROVED VERSION")
    print("="*70)
    
    try:
        plot_rmsd_analysis()
        create_2d_fel()
        plot_contact_analysis()
        plot_rmsf_analysis()
        create_summary_report()
        
        print("\n" + "="*70)
        print("✓ ANALYSIS COMPLETE!")
        print("="*70)
        print("\nAll visualizations have been saved to the current directory.")
        print("Check MD_Analysis_Summary.txt for detailed statistics.\n")
        
    except KeyboardInterrupt:
        print("\n\n⚠ Analysis interrupted by user")
        sys.exit(1)
    except Exception as e:
        print(f"\n\n✗ Fatal error: {str(e)}")
        import traceback
        traceback.print_exc()
        sys.exit(1)
```
