## Supplemental raw data for "A Computational Foundation Toward Targeting the ELMO1/DOCK2 Complex": PPI.py

```
#!/usr/bin/env python3
"""
Schrödinger Desmond MD Simulation Analysis Tool
================================================
Parses .eaf and .dat output files from Schrödinger Desmond simulations.
Generates RMSD/RMSF plots, 2D/3D Free Energy Landscapes, and protein-protein
interface binding-site analysis figures.

Usage
-----
  python desmond_analysis.py -eaf job-out_pl_1.eaf [-dat raw-data/] [-o results/] [-T 300]
  python desmond_analysis.py -eaf rep1.eaf rep2.eaf rep3.eaf -o results/

Requirements
------------
  pip install numpy matplotlib scipy seaborn scikit-learn
"""

import os
import re
import sys
import ast
import glob
import argparse
import warnings
import textwrap
import numpy as np
warnings.filterwarnings('ignore')

import matplotlib
matplotlib.use('Agg')
import matplotlib.pyplot as plt
import matplotlib.gridspec as gridspec
import matplotlib.ticker as ticker
from matplotlib.patches import Patch
from mpl_toolkits.mplot3d import Axes3D          # noqa: F401
from scipy.ndimage import gaussian_filter

try:
    import seaborn as sns
    sns.set_theme(style='ticks', palette='muted', font_scale=1.1)
    HAS_SNS = True
except ImportError:
    HAS_SNS = False

try:
    from sklearn.decomposition import PCA
    HAS_PCA = True
except ImportError:
    HAS_PCA = False

# ── Physical constants ─────────────────────────────────────────────────────────
R_KCAL = 1.987e-3   # kcal mol⁻¹ K⁻¹
R_KJ   = 8.314e-3   # kJ  mol⁻¹ K⁻¹

# ── Thresholds ─────────────────────────────────────────────────────────────────
HOTSPOT_FREQ  = 0.70   # ≥70 % frames → hotspot
IFACE_FREQ    = 0.30   # ≥30 % frames → interface

# Colour palette (colour-blind safe)
PALETTE = ['#2C7BB6', '#D7191C', '#1A9641', '#FDAE61', '#756BB1', '#31A354']
ITYPE_COLORS = {
    'hbond_bb':    '#3182BD',
    'hbond_ss':    '#6BAED6',
    'hbond_sb':    '#9ECAE1',
    'hbond_bs':    '#C6DBEF',
    'salt-bridge': '#E6550D',
    'pi-pi':       '#31A354',
    'pi-cat':      '#756BB1',
}
ITYPE_LABELS = {
    'hbond_bb':    'H-bond (bb–bb)',
    'hbond_ss':    'H-bond (sc–sc)',
    'hbond_sb':    'H-bond (sc–bb)',
    'hbond_bs':    'H-bond (bb–sc)',
    'salt-bridge': 'Salt bridge',
    'pi-pi':       'π–π stacking',
    'pi-cat':      'π–cation',
}

# ══════════════════════════════════════════════════════════════════════════════
# EAF PARSER
# ══════════════════════════════════════════════════════════════════════════════

class EAFSection:
    """One section extracted from a .eaf file."""
    __slots__ = ('stype', 'data')

    def __init__(self, stype: str, data: dict):
        self.stype = stype
        self.data  = data

    def result_array(self, key: str = 'Result') -> np.ndarray:
        v = self.data.get(key)
        if v is None:
            return np.array([])
        return np.asarray(v, dtype=float)

    def string_array(self, key: str = 'ProteinResidues') -> list:
        return self.data.get(key, [])

    def __repr__(self) -> str:
        return f"EAFSection(type={self.stype!r}, keys={list(self.data.keys())})"


def _parse_eaf_kv(line: str, data: dict) -> None:
    """Parse one 'Key = Value' line into data dict."""
    eq = line.find('=')
    if eq < 0:
        return
    key = line[:eq].strip()
    val = line[eq + 1:].strip()
    if not key:
        return

    # Float array: [0.0 1.2 ...]
    m = re.match(r'^\[([\d\s.eE+\-]+)\]$', val)
    if m:
        try:
            data[key] = [float(x) for x in m.group(1).split()]
            return
        except ValueError:
            pass

    # General bracket content
    m = re.match(r'^\[(.+)\]$', val, re.DOTALL)
    if m:
        inner = m.group(1).strip()
        quoted = re.findall(r'"([^"]*)"', inner)
        if quoted:
            data[key] = quoted
            return
        try:
            data[key] = [float(x) for x in inner.split()]
            return
        except ValueError:
            data[key] = inner
        return

    # Double-quoted string (may contain Python repr for ResultSummary / DictTag2ca)
    m = re.match(r'^"(.*)"$', val, re.DOTALL)
    if m:
        inner = m.group(1)
        if inner and (inner[0] in ('{', '[')):
            try:
                data[key] = ast.literal_eval(inner)
                return
            except Exception:
                pass
        data[key] = inner
        return

    # Bare integer
    try:
        data[key] = int(val)
        return
    except ValueError:
        pass
    # Bare float
    try:
        data[key] = float(val)
        return
    except ValueError:
        pass

    data[key] = val.strip('"\'')


def parse_eaf(filepath: str) -> list:
    """Parse a Schrödinger .eaf file → list of EAFSection."""
    sections = []
    current_type = None
    current_data: dict = {}
    depth = 0

    with open(filepath, 'r', errors='replace') as fh:
        for raw in fh:
            line = raw.strip()
            d_open  = line.count('{')
            d_close = line.count('}')

            # Detect new top-level section header (appears at depth==0)
            # Format: {SECTION_TYPE = {
            if depth == 0 and d_open >= 1:
                m = re.match(r'\{(\w+)\s*=\s*\{', line)
                if m:
                    current_type = m.group(1)
                    current_data = {}

            depth += d_open - d_close

            # Section inner-dict closed: depth dropped to 1 (first } of the pair)
            # At this point current_data is fully populated.
            if depth == 1 and current_type is not None:
                sections.append(EAFSection(current_type, current_data))
                current_type = None
                current_data = {}
                continue

            # Parse key-value pairs inside a section (depth == 2)
            if depth == 2 and '=' in line and current_type is not None:
                _parse_eaf_kv(line, current_data)

    return sections


# ══════════════════════════════════════════════════════════════════════════════
# DAT PARSERS
# ══════════════════════════════════════════════════════════════════════════════

def parse_dat_rmsd(filepath: str) -> dict:
    """Parse *_RMSD.dat → dict of column_name → np.ndarray."""
    header, rows = [], []
    with open(filepath) as fh:
        for line in fh:
            line = line.strip()
            if not line:
                continue
            if line.startswith('#'):
                header = line.lstrip('# ').split()
            else:
                try:
                    rows.append([float(x) for x in line.split()])
                except ValueError:
                    continue
    if not rows or not header:
        return {}
    arr = np.array(rows)
    return {h: arr[:, i] for i, h in enumerate(header) if i < arr.shape[1]}


def parse_dat_rmsf(filepath: str) -> dict:
    """Parse *_RMSF.dat → dict of column_name → list/array."""
    header, rows = [], []
    with open(filepath) as fh:
        for line in fh:
            line = line.strip()
            if not line:
                continue
            if line.startswith('#'):
                header = line.lstrip('# ').split()
            else:
                rows.append(line.split())
    if not rows or not header:
        return {}
    data: dict = {h: [] for h in header}
    for row in rows:
        for i, h in enumerate(header):
            if i < len(row):
                try:
                    data[h].append(float(row[i]))
                except ValueError:
                    data[h].append(row[i])
    for k in list(data.keys()):
        try:
            data[k] = np.array(data[k], dtype=float)
        except (ValueError, TypeError):
            pass
    return data


def parse_dat_contacts(filepath: str) -> dict:
    """Parse PL-Contacts_*.dat → {frame: count}."""
    frame_counts: dict = {}
    with open(filepath) as fh:
        for line in fh:
            line = line.strip()
            if not line or line.startswith('#'):
                continue
            try:
                frame = int(line.split()[0])
                frame_counts[frame] = frame_counts.get(frame, 0) + 1
            except (ValueError, IndexError):
                continue
    return frame_counts


# ══════════════════════════════════════════════════════════════════════════════
# FREE ENERGY LANDSCAPE
# ══════════════════════════════════════════════════════════════════════════════

def compute_fel_2d(cv1: np.ndarray, cv2: np.ndarray,
                   bins: int = 60, sigma: float = 1.5,
                   kT: float = R_KCAL * 300.0):
    """
    2D Free Energy Landscape: G = −kT ln[P(x,y)/P_max], normalised to 0 at minimum.
    Returns (X_grid, Y_grid, G_grid) with G in kcal/mol.
    """
    H, xe, ye = np.histogram2d(cv1, cv2, bins=bins)
    H = gaussian_filter(H.T.astype(float), sigma=sigma)
    H[H == 0] = H[H > 0].min() * 1e-3
    G = -kT * np.log(H / H.max())
    G -= G.min()
    xc = (xe[:-1] + xe[1:]) / 2
    yc = (ye[:-1] + ye[1:]) / 2
    X, Y = np.meshgrid(xc, yc)
    return X, Y, G


def compute_fel_1d(cv: np.ndarray, bins: int = 60, sigma: float = 1.5,
                   kT: float = R_KCAL * 300.0):
    """1D FEL from a single CV. Returns (x_centres, G)."""
    counts, edges = np.histogram(cv, bins=bins)
    counts = gaussian_filter(counts.astype(float), sigma=sigma)
    counts[counts == 0] = counts[counts > 0].min() * 1e-3
    G = -kT * np.log(counts / counts.max())
    G -= G.min()
    return (edges[:-1] + edges[1:]) / 2, G


# ══════════════════════════════════════════════════════════════════════════════
# PPI BINDING SITE ANALYSIS
# ══════════════════════════════════════════════════════════════════════════════

def _chain_of(res: str) -> str:
    """Extract chain letter from 'A:GLY_1' → 'A'."""
    return res.split(':')[0]


def analyse_ppi(ppi_summary: dict, n_frames: int) -> dict:
    """
    Analyse ProtProtInter ResultSummary dict from the EAF file.

    Parameters
    ----------
    ppi_summary : {itype: {(res1, res2): count, ...}, ...}
    n_frames    : total MD frames

    Returns
    -------
    dict with keys:
      score_A, score_B          – max inter-chain contact freq per residue
      cum_score_A, cum_score_B  – cumulative (sum) contact freq per residue
      hotspot_A/B, iface_A/B   – classified residue lists
      by_type                   – {itype: {res: freq}} for inter-chain contacts
      inter_pairs               – {(rA, rB): {itype: freq}}
      all_chains                – set of chain IDs found
    """
    chains = set()
    inter_pairs: dict = {}
    by_type: dict = {}
    max_score: dict = {'A_side': {}, 'B_side': {}}
    cum_score: dict = {'A_side': {}, 'B_side': {}}

    for itype, pairs in ppi_summary.items():
        for (r1, r2), count in pairs.items():
            c1, c2 = _chain_of(r1), _chain_of(r2)
            chains.update([c1, c2])
            if c1 == c2:
                continue   # intra-chain — skip for interface analysis
            freq = count / n_frames
            # Canonical ordering: A_side = the first (lexicographically lower) chain
            if c1 < c2:
                rA, rB = r1, r2
            else:
                rA, rB = r2, r1
            inter_pairs.setdefault((rA, rB), {})[itype] = freq
            by_type.setdefault(itype, {}).setdefault(rA, 0)
            by_type[itype][rA] = by_type[itype].get(rA, 0) + freq
            by_type[itype].setdefault(rB, 0)
            by_type[itype][rB] = by_type[itype].get(rB, 0) + freq
            # Max score
            max_score['A_side'][rA] = max(max_score['A_side'].get(rA, 0), freq)
            max_score['B_side'][rB] = max(max_score['B_side'].get(rB, 0), freq)
            # Cumulative score (sum of all inter-chain contacts involving this residue)
            cum_score['A_side'][rA] = cum_score['A_side'].get(rA, 0) + freq
            cum_score['B_side'][rB] = cum_score['B_side'].get(rB, 0) + freq

    sA = max_score['A_side']
    sB = max_score['B_side']
    hotspot_A = sorted([r for r, f in sA.items() if f >= HOTSPOT_FREQ], key=lambda r: -sA[r])
    iface_A   = sorted([r for r, f in sA.items() if IFACE_FREQ <= f < HOTSPOT_FREQ], key=lambda r: -sA[r])
    hotspot_B = sorted([r for r, f in sB.items() if f >= HOTSPOT_FREQ], key=lambda r: -sB[r])
    iface_B   = sorted([r for r, f in sB.items() if IFACE_FREQ <= f < HOTSPOT_FREQ], key=lambda r: -sB[r])

    return {
        'score_A':     sA,
        'score_B':     sB,
        'cum_score_A': cum_score['A_side'],
        'cum_score_B': cum_score['B_side'],
        'hotspot_A':   hotspot_A,
        'iface_A':     iface_A,
        'hotspot_B':   hotspot_B,
        'iface_B':     iface_B,
        'by_type':     by_type,
        'inter_pairs': inter_pairs,
        'all_chains':  sorted(chains),
    }


# ══════════════════════════════════════════════════════════════════════════════
# MAIN ANALYSER CLASS
# ══════════════════════════════════════════════════════════════════════════════

class DesmondAnalysis:
    """Orchestrates loading, analysis, and figure generation."""

    def __init__(self, eaf_files: list, dat_dir: str = None,
                 output_dir: str = 'desmond_analysis_output',
                 temperature: float = 300.0, label: str = None):
        self.eaf_files   = eaf_files
        self.dat_dir     = dat_dir
        self.output_dir  = output_dir
        self.temperature = temperature
        self.kT          = R_KCAL * temperature
        self.label       = label or os.path.basename(eaf_files[0]).replace('.eaf', '') if eaf_files else 'MD'

        os.makedirs(output_dir, exist_ok=True)

        # Parsed data containers
        self.rmsd_data:  dict = {}   # key → np.ndarray (per frame)
        self.rmsf_data:  dict = {}   # 'residues' → list[str], 'values' → array
        self.ppi_result: dict = {}
        self.n_frames:   int  = 0
        self.contacts_ts: np.ndarray = np.array([])  # contacts per frame
        self.dat_rmsd:   dict = {}
        self.dat_rmsf:   dict = {}
        # New containers for the 4 additional analyses
        self.rg_data:    np.ndarray = np.array([])   # Rg per frame
        self.ss_matrix:  np.ndarray = None           # (n_res, n_frames) SS codes
        self.ss_residues: list = []
        self.contact_persist:        np.ndarray = None   # (n_res, n_frames) binary
        self.contact_persist_labels: list = []
        self.unbound_rmsf: dict = {}                 # for differential RMSF

    # ── Data loading ─────────────────────────────────────────────────────────

    def load(self):
        """Load all available data sources."""
        self._load_eaf()
        self._load_dat()
        print(f"\n  Frames : {self.n_frames}")
        print(f"  RMSD series available: {list(self.rmsd_data.keys())}")
        if self.ppi_result:
            print(f"  Inter-chain pairs: {self.ppi_result.get('n_inter_pairs', '?')}")
            print(f"  Hotspot A: {self.ppi_result.get('hotspot_A', [])}")
            print(f"  Hotspot B: {self.ppi_result.get('hotspot_B', [])}")

    def _load_eaf(self):
        for path in self.eaf_files:
            print(f"\nParsing EAF: {path}")
            sections = parse_eaf(path)
            print(f"  Found {len(sections)} sections: "
                  f"{sorted(set(s.stype for s in sections))}")
            self._extract_from_sections(sections)

    def _extract_from_sections(self, sections: list):
        # -- RMSD --
        rmsd_secs = [s for s in sections if s.stype == 'RMSD']
        for s in rmsd_secs:
            sel = s.data.get('SelectionType', 'Unknown')
            arr = s.result_array('Result')
            if arr.size:
                key = sel
                if key not in self.rmsd_data:
                    self.rmsd_data[key] = arr
                else:
                    self.rmsd_data[key] = np.concatenate([self.rmsd_data[key], arr])
                self.n_frames = max(self.n_frames, len(arr))

        # -- RMSF --
        rmsf_secs = [s for s in sections if s.stype == 'RMSF']
        for s in rmsf_secs:
            sel = s.data.get('SelectionType', 'Unknown')
            arr = s.result_array('Result')
            res = s.string_array('ProteinResidues')
            if arr.size and res and 'CA' in sel:
                self.rmsf_data = {'residues': res, 'values': arr,
                                  'selection': sel}
                break
        # If no CA, take first RMSF with residues
        if not self.rmsf_data:
            for s in rmsf_secs:
                arr = s.result_array('Result')
                res = s.string_array('ProteinResidues')
                if arr.size and res:
                    self.rmsf_data = {'residues': res, 'values': arr,
                                      'selection': s.data.get('SelectionType', '')}
                    break

        # -- ProtProtInter / ProtLigInter --
        ppi_secs = [s for s in sections
                    if s.stype in ('ProtProtInter', 'ProtLigInter')]
        for s in ppi_secs:
            summary = s.data.get('ResultSummary')
            if summary and isinstance(summary, dict):
                nf = self.n_frames if self.n_frames else 1
                self.ppi_result = analyse_ppi(summary, nf)
                self.ppi_result['n_inter_pairs'] = len(self.ppi_result['inter_pairs'])
                break

        # -- Rad_Gyration --
        rg_secs = [s for s in sections if s.stype == 'Rad_Gyration']
        for s in rg_secs:
            arr = s.result_array('Result')
            if arr.size and self.rg_data.size == 0:
                self.rg_data = arr
                break

        # -- SecondaryStructure --
        ss_secs = [s for s in sections if s.stype == 'SecondaryStructure']
        for s in ss_secs:
            res = s.string_array('ProteinResidues')
            results = s.data.get('Result', [])
            if res and results and isinstance(results, list) and self.ss_matrix is None:
                try:
                    # Each element is a dot-separated string of int SS codes, one per residue
                    matrix = np.array(
                        [[int(c) for c in row.split('.')]
                         for row in results if isinstance(row, str)],
                        dtype=np.int8
                    ).T    # → (n_residues, n_frames)
                    self.ss_matrix   = matrix
                    self.ss_residues = res
                    print(f"  SecondaryStructure: {matrix.shape[0]} residues × "
                          f"{matrix.shape[1]} frames")
                except (ValueError, IndexError) as exc:
                    print(f"  SecondaryStructure parse error: {exc}")
            break

    def _load_dat(self):
        if not self.dat_dir or not os.path.isdir(self.dat_dir):
            return
        print(f"\nLoading DAT files from: {self.dat_dir}")
        dat_files = sorted(glob.glob(os.path.join(self.dat_dir, '*.dat')))
        # Prefer P_RMSD / P_RMSF (protein) over L_* (ligand)
        dat_files.sort(key=lambda f: (0 if os.path.basename(f).startswith('P') else 1))
        for f in dat_files:
            name = os.path.basename(f)
            if 'RMSD' in name.upper() and not self.dat_rmsd:
                self.dat_rmsd = parse_dat_rmsd(f)
                print(f"  RMSD dat: {name} → {list(self.dat_rmsd.keys())}")
            elif 'RMSF' in name.upper() and not self.dat_rmsf:
                self.dat_rmsf = parse_dat_rmsf(f)
                print(f"  RMSF dat: {name} → {list(self.dat_rmsf.keys())}")
            elif 'Contact' in name or 'contact' in name:
                counts = parse_dat_contacts(f)
                if counts:
                    max_frame = max(counts.keys())
                    ts = np.zeros(max_frame + 1)
                    for fr, c in counts.items():
                        ts[fr] = c
                    if self.contacts_ts.size == 0:
                        self.contacts_ts = ts
                    else:
                        n = min(len(self.contacts_ts), len(ts))
                        self.contacts_ts = self.contacts_ts[:n] + ts[:n]
            elif 'Properties' in name or 'properties' in name:
                if self.rg_data.size == 0:
                    prop = parse_dat_rmsd(f)
                    if 'rGyr' in prop:
                        self.rg_data = prop['rGyr']
                        print(f"  Rg data from: {name} ({len(self.rg_data)} frames)")

        self._build_contact_persistence()

    def _build_contact_persistence(self):
        """Build binary (n_res, n_frames) contact matrix from PL-Contacts_*.dat.

        Schrödinger contact DAT format (columns):
          Frame#  Residue#  Chain  ResName  AtomName  LigandFragment  LigandAtomName
        Residue label is reconstructed as 'Chain:ResName_Residue#'.
        """
        if not self.dat_dir or not os.path.isdir(self.dat_dir):
            return
        contact_files = sorted(
            glob.glob(os.path.join(self.dat_dir, '*Contact*.dat')) +
            glob.glob(os.path.join(self.dat_dir, '*contact*.dat'))
        )
        if not contact_files:
            return

        observations: set = set()

        for fpath in contact_files:
            with open(fpath, errors='replace') as fh:
                for raw in fh:
                    line = raw.strip()
                    if not line or line.startswith('#'):
                        continue
                    parts = line.split()
                    if len(parts) < 4:
                        continue
                    try:
                        frame   = int(parts[0])
                        resnum  = int(parts[1])
                        chain   = parts[2]
                        resname = parts[3]
                    except (ValueError, IndexError):
                        continue
                    res_label = f"{chain}:{resname}_{resnum}"
                    observations.add((frame, res_label))

        if not observations:
            return

        all_frames = sorted(set(f for f, _ in observations))
        all_res    = sorted(set(r for _, r in observations), key=_resid_sort_key)

        res_idx   = {r: i for i, r in enumerate(all_res)}
        frame_idx = {f: i for i, f in enumerate(all_frames)}

        mat = np.zeros((len(all_res), len(all_frames)), dtype=np.int8)
        for frame, res in observations:
            mat[res_idx[res], frame_idx[frame]] = 1

        self.contact_persist        = mat
        self.contact_persist_labels = all_res
        print(f"  Contact persistence matrix: "
              f"{len(all_res)} residues × {len(all_frames)} frames")

    def load_unbound_rmsf(self, unbound_eaf_path: str):
        """Parse an unbound simulation EAF and extract RMSF for differential analysis."""
        if not unbound_eaf_path or not os.path.isfile(unbound_eaf_path):
            print(f"  Unbound EAF not found: {unbound_eaf_path}")
            return
        print(f"\nLoading unbound RMSF from: {unbound_eaf_path}")
        sections  = parse_eaf(unbound_eaf_path)
        rmsf_secs = [s for s in sections if s.stype == 'RMSF']
        for s in rmsf_secs:
            sel = s.data.get('SelectionType', '')
            arr = s.result_array('Result')
            res = s.string_array('ProteinResidues')
            if arr.size and res and 'CA' in sel:
                self.unbound_rmsf = {'residues': res, 'values': arr}
                print(f"  Unbound RMSF (CA): {len(res)} residues")
                return
        for s in rmsf_secs:
            arr = s.result_array('Result')
            res = s.string_array('ProteinResidues')
            if arr.size and res:
                self.unbound_rmsf = {'residues': res, 'values': arr}
                print(f"  Unbound RMSF: {len(res)} residues")
                return
        print("  No RMSF section found in unbound EAF.")

    # ── Helpers ───────────────────────────────────────────────────────────────

    def _savefig(self, fig, name: str, dpi: int = 300):
        path = os.path.join(self.output_dir, name)
        fig.savefig(path, dpi=dpi, bbox_inches='tight', facecolor='white')
        print(f"  Saved → {path}")
        plt.close(fig)

    def _primary_rmsd(self) -> tuple:
        """Return (label, array) for the most useful RMSD series."""
        preferred = ['Backbone', 'CA', 'All_Heavy', 'Unknown']
        for p in preferred:
            if p in self.rmsd_data:
                return p, self.rmsd_data[p]
        if self.rmsd_data:
            k = next(iter(self.rmsd_data))
            return k, self.rmsd_data[k]
        # Fall back to dat file
        for key in ('Prot_Backbone', 'Prot_CA', 'Prot_All_Heavy'):
            if key in self.dat_rmsd:
                return key, self.dat_rmsd[key]
        return None, np.array([])

    @staticmethod
    def _running_avg(arr: np.ndarray, window: int) -> np.ndarray:
        w = max(1, min(window, len(arr)))
        return np.convolve(arr, np.ones(w) / w, mode='same')

    # ── Plot 01 – RMSD ───────────────────────────────────────────────────────

    def plot_rmsd(self):
        # Collect series from eaf and dat
        series: dict = {}
        for k, v in self.rmsd_data.items():
            series[k] = v
        for k in ('Prot_CA', 'Prot_Backbone', 'Lig_wrt_Protein', 'Lig_wrt_Ligand'):
            if k in self.dat_rmsd and k not in series:
                series[k] = self.dat_rmsd[k]

        if not series:
            print("  No RMSD data found – skipping.")
            return

        fig, axes = plt.subplots(2, 1, figsize=(12, 7),
                                 gridspec_kw={'height_ratios': [3, 1]})
        ax, ax_dist = axes

        colors = PALETTE
        for i, (lbl, arr) in enumerate(series.items()):
            frames = np.arange(len(arr))
            color  = colors[i % len(colors)]
            ax.plot(frames, arr, color=color, lw=0.8, alpha=0.5)
            window = max(5, len(arr) // 50)
            avg = self._running_avg(arr, window)
            ax.plot(frames, avg, color=color, lw=2.2,
                    label=f"{lbl}  (mean={arr.mean():.2f} Å)")

        ax.set_xlabel('Frame', fontsize=13)
        ax.set_ylabel('RMSD (Å)', fontsize=13)
        ax.set_title(f'RMSD – {self.label}', fontsize=14, fontweight='bold')
        ax.legend(fontsize=10, framealpha=0.8)
        ax.grid(True, alpha=0.25)

        # Distribution (primary series)
        _, primary = self._primary_rmsd()
        if primary.size:
            ax_dist.hist(primary, bins=50, color=colors[0], alpha=0.75,
                         edgecolor='white', orientation='horizontal')
            ax_dist.axhline(primary.mean(), color='red', ls='--', lw=1.5,
                            label=f'Mean = {primary.mean():.2f} Å')
            ax_dist.set_xlabel('Count', fontsize=11)
            ax_dist.set_ylabel('RMSD (Å)', fontsize=11)
            ax_dist.legend(fontsize=9)
            ax_dist.grid(True, alpha=0.25)

        plt.tight_layout()
        self._savefig(fig, '01_RMSD.png')

    # ── Plot 02 – RMSF ───────────────────────────────────────────────────────

    def plot_rmsf(self):
        if not self.rmsf_data:
            # Try DAT
            if self.dat_rmsf:
                self._plot_rmsf_from_dat()
            else:
                print("  No RMSF data – skipping.")
            return

        residues = self.rmsf_data['residues']
        values   = self.rmsf_data['values']

        # Parse residue numbers and chains
        resids, chains = [], []
        for r in residues:
            m = re.match(r'([A-Za-z]):\w+_(\d+)', r)
            if m:
                chains.append(m.group(1))
                resids.append(int(m.group(2)))
            else:
                chains.append('A')
                resids.append(len(resids))

        resids = np.array(resids)
        chains = np.array(chains)
        unique_chains = sorted(set(chains))

        fig, ax = plt.subplots(figsize=(14, 5))
        for i, ch in enumerate(unique_chains):
            mask = chains == ch
            bar_colors = np.where(
                values[mask] > values.mean() + 2 * values.std(),
                PALETTE[i] if i < len(PALETTE) else 'red',
                _lighten(PALETTE[i % len(PALETTE)], 0.4)
            )
            ax.bar(resids[mask], values[mask], color=bar_colors,
                   width=1.0, alpha=0.85, label=f'Chain {ch}')

        # Threshold line
        thr = values.mean() + 2 * values.std()
        ax.axhline(thr, color='red', ls='--', lw=1.5, alpha=0.8,
                   label=f'Mean+2σ = {thr:.2f} Å')

        # Annotate hotspot residues on the RMSF plot
        if self.ppi_result:
            all_hot = (list(self.ppi_result.get('hotspot_A', [])) +
                       list(self.ppi_result.get('hotspot_B', [])))
            for r in all_hot:
                m = re.match(r'\w+:(\w+)_(\d+)', r)
                if m:
                    rid = int(m.group(2))
                    idx_arr = np.where(resids == rid)[0]
                    if idx_arr.size:
                        ax.annotate('★', xy=(rid, values[idx_arr[0]]),
                                    ha='center', va='bottom',
                                    fontsize=8, color='gold', fontweight='bold')

        ax.set_xlabel('Residue Number', fontsize=13)
        ax.set_ylabel('RMSF (Å)', fontsize=13)
        ax.set_title(f'Per-Residue RMSF – {self.label}', fontsize=14, fontweight='bold')
        legend_handles = [Patch(color=PALETTE[i % len(PALETTE)], label=f'Chain {ch}')
                          for i, ch in enumerate(unique_chains)]
        legend_handles.append(plt.Line2D([0], [0], color='red', ls='--',
                                         label=f'Mean+2σ = {thr:.2f} Å'))
        if self.ppi_result:
            legend_handles.append(plt.Line2D([0], [0], marker='*', color='w',
                                             markerfacecolor='gold', markersize=12,
                                             label='Hotspot residue'))
        ax.legend(handles=legend_handles, fontsize=10, framealpha=0.8)
        ax.grid(True, alpha=0.2, axis='y')
        plt.tight_layout()
        self._savefig(fig, '02_RMSF.png')

    def _plot_rmsf_from_dat(self):
        d = self.dat_rmsf
        key = 'CA' if 'CA' in d else 'Backbone' if 'Backbone' in d else None
        if key is None:
            return
        fig, ax = plt.subplots(figsize=(12, 4))
        vals = np.asarray(d[key])
        ax.bar(range(len(vals)), vals, color=PALETTE[0], alpha=0.8, width=1.0)
        ax.set_xlabel('Residue index', fontsize=12)
        ax.set_ylabel('RMSF (Å)', fontsize=12)
        ax.set_title(f'RMSF ({key}) – {self.label}', fontweight='bold')
        ax.grid(True, alpha=0.2, axis='y')
        plt.tight_layout()
        self._savefig(fig, '02_RMSF.png')

    # ── Plot 03 – 2D FEL ─────────────────────────────────────────────────────

    def plot_fel_2d(self):
        cv1_label, cv1 = self._cv1()
        cv2_label, cv2 = self._cv2()
        if cv1 is None or cv2 is None:
            print("  Insufficient CVs for 2D FEL – skipping.")
            return
        n = min(len(cv1), len(cv2))
        if n < 20:
            return
        cv1, cv2 = cv1[:n], cv2[:n]

        X, Y, G = compute_fel_2d(cv1, cv2, bins=60, sigma=1.5, kT=self.kT)
        min_idx = np.unravel_index(G.argmin(), G.shape)
        n_levels = 15
        G_max_plot = min(G.max(), 5.0)  # Cap at 5 kcal/mol for colour scale

        fig, (ax2d, ax1d) = plt.subplots(1, 2, figsize=(13, 5))

        # 2D contour
        cf = ax2d.contourf(X, Y, G, levels=n_levels,
                           vmin=0, vmax=G_max_plot, cmap='RdYlBu_r', extend='max')
        ax2d.contour(X, Y, G, levels=8, colors='black', alpha=0.35, linewidths=0.6)
        cb = plt.colorbar(cf, ax=ax2d, pad=0.02)
        cb.set_label('ΔG (kcal/mol)', fontsize=11)
        ax2d.plot(X[min_idx], Y[min_idx], 'w*', markersize=18, zorder=5,
                  label=f'Global min ({X[min_idx]:.2f}, {Y[min_idx]:.2f})')
        ax2d.set_xlabel(cv1_label, fontsize=12)
        ax2d.set_ylabel(cv2_label, fontsize=12)
        ax2d.set_title('2D Free Energy Landscape', fontsize=13, fontweight='bold')
        ax2d.legend(fontsize=9, framealpha=0.8)

        # 1D projections on the right
        x1d_1, G1d_1 = compute_fel_1d(cv1, kT=self.kT)
        x1d_2, G1d_2 = compute_fel_1d(cv2, kT=self.kT)
        ax1d.plot(x1d_1, G1d_1, color=PALETTE[0], lw=2.0, label=cv1_label)
        ax1d_r = ax1d.twinx()
        ax1d_r.plot(x1d_2, G1d_2, color=PALETTE[1], lw=2.0, ls='--', label=cv2_label)
        ax1d.set_xlabel('CV value (Å)', fontsize=12)
        ax1d.set_ylabel(f'ΔG [{cv1_label}] (kcal/mol)', fontsize=11, color=PALETTE[0])
        ax1d_r.set_ylabel(f'ΔG [{cv2_label}] (kcal/mol)', fontsize=11, color=PALETTE[1])
        ax1d.set_title('1D FEL Projections', fontsize=13, fontweight='bold')
        lines1, lab1 = ax1d.get_legend_handles_labels()
        lines2, lab2 = ax1d_r.get_legend_handles_labels()
        ax1d.legend(lines1 + lines2, lab1 + lab2, fontsize=9)
        ax1d.grid(True, alpha=0.25)

        plt.suptitle(f'Free Energy Landscape – {self.label}', fontsize=14,
                     fontweight='bold', y=1.01)
        plt.tight_layout()
        self._savefig(fig, '03_FEL_2D.png')

    # ── Plot 04 – 3D FEL ─────────────────────────────────────────────────────

    def plot_fel_3d(self):
        cv1_label, cv1 = self._cv1()
        cv2_label, cv2 = self._cv2()
        if cv1 is None or cv2 is None:
            return
        n = min(len(cv1), len(cv2))
        if n < 20:
            return
        X, Y, G = compute_fel_2d(cv1[:n], cv2[:n], bins=50, sigma=1.5, kT=self.kT)
        G_plot = np.clip(G, 0, 5.0)

        fig = plt.figure(figsize=(13, 5))

        ax3d = fig.add_subplot(121, projection='3d')
        surf = ax3d.plot_surface(X, Y, G_plot, cmap='RdYlBu_r',
                                 linewidth=0, antialiased=True, alpha=0.92)
        fig.colorbar(surf, ax=ax3d, shrink=0.55, aspect=12,
                     label='ΔG (kcal/mol)')
        ax3d.set_xlabel(cv1_label, fontsize=9, labelpad=6)
        ax3d.set_ylabel(cv2_label, fontsize=9, labelpad=6)
        ax3d.set_zlabel('ΔG (kcal/mol)', fontsize=9, labelpad=6)
        ax3d.set_title('3D FEL Surface', fontsize=12, fontweight='bold')
        ax3d.view_init(elev=30, azim=-60)

        # Overhead contour (same data)
        ax2 = fig.add_subplot(122)
        cf = ax2.contourf(X, Y, G_plot, levels=20, cmap='RdYlBu_r', vmin=0, vmax=5)
        ax2.contour(X, Y, G_plot, levels=10, colors='black', alpha=0.3, linewidths=0.5)
        plt.colorbar(cf, ax=ax2, label='ΔG (kcal/mol)')
        min_idx = np.unravel_index(G_plot.argmin(), G_plot.shape)
        ax2.plot(X[min_idx], Y[min_idx], 'w*', markersize=16, zorder=5)
        ax2.set_xlabel(cv1_label, fontsize=12)
        ax2.set_ylabel(cv2_label, fontsize=12)
        ax2.set_title('Overhead View', fontsize=12, fontweight='bold')

        plt.suptitle(f'3D Free Energy Landscape – {self.label}', fontsize=14,
                     fontweight='bold')
        plt.tight_layout()
        self._savefig(fig, '04_FEL_3D.png')

    # ── Plot 05 – PPI Contact Map ─────────────────────────────────────────────

    def plot_ppi_contact_map(self):
        if not self.ppi_result or not self.ppi_result.get('inter_pairs'):
            print("  No PPI inter-chain contacts – skipping contact map.")
            return

        pairs   = self.ppi_result['inter_pairs']
        score_A = self.ppi_result['score_A']
        score_B = self.ppi_result['score_B']

        # Build sorted residue lists
        all_A = sorted(score_A.keys(), key=_resid_sort_key)
        all_B = sorted(score_B.keys(), key=_resid_sort_key)

        idx_A = {r: i for i, r in enumerate(all_A)}
        idx_B = {r: i for i, r in enumerate(all_B)}

        mat = np.zeros((len(all_B), len(all_A)))
        for (rA, rB), itypes in pairs.items():
            if rA in idx_A and rB in idx_B:
                ia, ib = idx_A[rA], idx_B[rB]
                mat[ib, ia] = max(itypes.values())   # max single-type freq

        fig, ax = plt.subplots(figsize=(max(8, len(all_A)//3), max(5, len(all_B)//3 + 2)))
        im = ax.imshow(mat, aspect='auto', cmap='hot_r', vmin=0, vmax=1,
                       origin='lower')
        cb = plt.colorbar(im, ax=ax, fraction=0.03)
        cb.set_label('Max contact frequency', fontsize=11)

        step_A = max(1, len(all_A) // 20)
        step_B = max(1, len(all_B) // 10)
        ax.set_xticks(range(0, len(all_A), step_A))
        ax.set_xticklabels([all_A[i].split(':')[1] for i in range(0, len(all_A), step_A)],
                           rotation=45, ha='right', fontsize=7)
        ax.set_yticks(range(0, len(all_B), step_B))
        ax.set_yticklabels([all_B[i].split(':')[1] for i in range(0, len(all_B), step_B)],
                           fontsize=8)

        chains = self.ppi_result.get('all_chains', ['A', 'B'])
        ax.set_xlabel(f'Chain {chains[0]} residues', fontsize=12)
        ax.set_ylabel(f'Chain {chains[1]} residues', fontsize=12)
        ax.set_title(f'Interface Contact Frequency Map – {self.label}',
                     fontsize=13, fontweight='bold')

        # Mark hotspots with cyan lines
        for r in self.ppi_result.get('hotspot_A', []):
            if r in idx_A:
                ax.axvline(idx_A[r], color='cyan', lw=1.2, alpha=0.7)
        for r in self.ppi_result.get('hotspot_B', []):
            if r in idx_B:
                ax.axhline(idx_B[r], color='cyan', lw=1.2, alpha=0.7)

        plt.tight_layout()
        self._savefig(fig, '05_PPI_ContactMap.png')

    # ── Plot 06 – Hotspot Bar Charts ─────────────────────────────────────────

    def plot_ppi_hotspots(self):
        if not self.ppi_result:
            print("  No PPI data – skipping hotspot plot.")
            return

        score_A = self.ppi_result['score_A']
        score_B = self.ppi_result['score_B']
        if not score_A and not score_B:
            return

        chains = self.ppi_result.get('all_chains', ['A', 'B'])
        datasets = []
        if score_A:
            datasets.append((chains[0], score_A))
        if score_B and len(chains) > 1:
            datasets.append((chains[1], score_B))

        fig, axes = plt.subplots(len(datasets), 1,
                                 figsize=(14, 4 * len(datasets)), squeeze=False)

        for ax, (ch, scores) in zip(axes[:, 0], datasets):
            sorted_res = sorted(scores.keys(), key=_resid_sort_key)
            sorted_freq = [scores[r] for r in sorted_res]
            bar_colors = [
                '#D7191C' if f >= HOTSPOT_FREQ
                else '#FDAE61' if f >= IFACE_FREQ
                else '#ABD9E9'
                for f in sorted_freq
            ]
            ax.bar(range(len(sorted_res)), sorted_freq, color=bar_colors, alpha=0.9)
            ax.axhline(HOTSPOT_FREQ, color='red', ls='--', lw=1.5,
                       label=f'Hotspot ≥{HOTSPOT_FREQ:.0%}')
            ax.axhline(IFACE_FREQ, color='orange', ls='--', lw=1.5,
                       label=f'Interface ≥{IFACE_FREQ:.0%}')

            # Label hotspot residues
            for i, (r, f) in enumerate(zip(sorted_res, sorted_freq)):
                if f >= HOTSPOT_FREQ:
                    ax.text(i, f + 0.01, r.split(':')[1], ha='center', va='bottom',
                            fontsize=7, rotation=90, color='darkred', fontweight='bold')

            step = max(1, len(sorted_res) // 20)
            ax.set_xticks(range(0, len(sorted_res), step))
            ax.set_xticklabels(
                [sorted_res[i].split(':')[1] for i in range(0, len(sorted_res), step)],
                rotation=45, ha='right', fontsize=8)
            ax.set_ylim(0, min(1.2, max(sorted_freq) * 1.3 + 0.1))
            ax.set_ylabel('Max contact frequency', fontsize=11)
            ax.set_title(f'Chain {ch} – Interface Residue Scores', fontsize=12,
                         fontweight='bold')
            legend_patches = [
                Patch(color='#D7191C', label=f'Hotspot (≥{HOTSPOT_FREQ:.0%})'),
                Patch(color='#FDAE61', label=f'Interface (≥{IFACE_FREQ:.0%})'),
                Patch(color='#ABD9E9', label='Peripheral'),
            ]
            ax.legend(handles=legend_patches, fontsize=9, loc='upper right')
            ax.grid(True, alpha=0.2, axis='y')

        plt.suptitle(f'Binding-Site Residue Scores – {self.label}',
                     fontsize=14, fontweight='bold', y=1.01)
        plt.tight_layout()
        self._savefig(fig, '06_PPI_Hotspots.png')

    # ── Plot 07 – Interaction Type Breakdown ─────────────────────────────────

    def plot_interaction_breakdown(self):
        if not self.ppi_result or not self.ppi_result.get('by_type'):
            print("  No PPI interaction-type data – skipping breakdown plot.")
            return

        by_type = self.ppi_result['by_type']
        n_frames = self.n_frames if self.n_frames else 1

        # Sum total contacts per interaction type (inter-chain only)
        inter_pairs = self.ppi_result.get('inter_pairs', {})
        type_totals: dict = {}
        for (rA, rB), itypes in inter_pairs.items():
            for itype, freq in itypes.items():
                type_totals[itype] = type_totals.get(itype, 0) + freq

        if not type_totals:
            return

        sorted_types = sorted(type_totals, key=lambda x: -type_totals[x])
        totals = [type_totals[t] for t in sorted_types]
        labels = [ITYPE_LABELS.get(t, t) for t in sorted_types]
        colors = [ITYPE_COLORS.get(t, '#888888') for t in sorted_types]

        fig, (ax_pie, ax_bar) = plt.subplots(1, 2, figsize=(13, 5))

        # Pie chart
        wedges, texts, autotexts = ax_pie.pie(
            totals, labels=labels, colors=colors, autopct='%1.1f%%',
            pctdistance=0.8, startangle=90,
            wedgeprops={'edgecolor': 'white', 'linewidth': 1.5})
        for t in autotexts:
            t.set_fontsize(9)
        ax_pie.set_title('Interaction Type Distribution\n(inter-chain, by cumulative frequency)',
                         fontsize=12, fontweight='bold')

        # Stacked bar: top-10 hotspot residues A
        all_res_A = sorted(self.ppi_result['score_A'],
                           key=lambda r: -self.ppi_result['score_A'][r])[:10]
        if all_res_A:
            bottom = np.zeros(len(all_res_A))
            for itype in sorted_types:
                vals = np.array([by_type.get(itype, {}).get(r, 0) for r in all_res_A])
                ax_bar.bar(range(len(all_res_A)), vals, bottom=bottom,
                           color=ITYPE_COLORS.get(itype, '#888'),
                           label=ITYPE_LABELS.get(itype, itype), alpha=0.9)
                bottom += vals
            chains = self.ppi_result.get('all_chains', ['A', 'B'])
            ax_bar.set_xticks(range(len(all_res_A)))
            ax_bar.set_xticklabels([r.split(':')[1] for r in all_res_A],
                                   rotation=45, ha='right', fontsize=9)
            ax_bar.set_ylabel('Cumulative contact frequency', fontsize=11)
            ax_bar.set_title(f'Top-10 Chain {chains[0]} Interface Residues\n(by interaction type)',
                             fontsize=12, fontweight='bold')
            ax_bar.legend(fontsize=8, bbox_to_anchor=(1.01, 1), loc='upper left')
            ax_bar.grid(True, alpha=0.2, axis='y')

        plt.suptitle(f'Interaction Breakdown – {self.label}',
                     fontsize=14, fontweight='bold')
        plt.tight_layout()
        self._savefig(fig, '07_Interaction_Breakdown.png')

    # ── Plot 08 – Summary Dashboard ──────────────────────────────────────────

    def plot_dashboard(self):
        fig = plt.figure(figsize=(18, 12))
        fig.suptitle(f'MD Simulation Analysis – {self.label}',
                     fontsize=16, fontweight='bold', y=1.01)
        gs_ = gridspec.GridSpec(3, 3, figure=fig, hspace=0.42, wspace=0.35)

        # ── Panel A: RMSD time series ──
        ax_rmsd = fig.add_subplot(gs_[0, :2])
        _, primary = self._primary_rmsd()
        if primary.size:
            frames = np.arange(len(primary))
            ax_rmsd.plot(frames, primary, color=PALETTE[0], lw=0.8, alpha=0.4)
            avg = self._running_avg(primary, max(5, len(primary) // 40))
            ax_rmsd.plot(frames, avg, color=PALETTE[0], lw=2.2)
            ax_rmsd.set_xlabel('Frame'); ax_rmsd.set_ylabel('RMSD (Å)')
            ax_rmsd.set_title('Backbone RMSD', fontweight='bold')
            ax_rmsd.grid(True, alpha=0.2)

        # ── Panel B: RMSD distribution ──
        ax_rdist = fig.add_subplot(gs_[0, 2])
        if primary.size:
            ax_rdist.hist(primary, bins=40, color=PALETTE[0], alpha=0.8,
                          edgecolor='white', orientation='horizontal')
            ax_rdist.axhline(primary.mean(), color='red', ls='--', lw=1.8,
                             label=f'{primary.mean():.2f} Å')
            ax_rdist.set_xlabel('Count'); ax_rdist.set_ylabel('RMSD (Å)')
            ax_rdist.set_title('RMSD Distribution', fontweight='bold')
            ax_rdist.legend(fontsize=8)
            ax_rdist.grid(True, alpha=0.2)

        # ── Panel C: RMSF ──
        ax_rmsf = fig.add_subplot(gs_[1, :2])
        if self.rmsf_data:
            residues = self.rmsf_data['residues']
            rmsf_vals = self.rmsf_data['values']
            resids_num = []
            for r in residues:
                m = re.match(r'[A-Z]:\w+_(\d+)', r)
                resids_num.append(int(m.group(1)) if m else len(resids_num))
            ax_rmsf.bar(resids_num, rmsf_vals, color=PALETTE[0], alpha=0.75, width=1.0)
            ax_rmsf.axhline(rmsf_vals.mean() + 2 * rmsf_vals.std(),
                            color='red', ls='--', lw=1.5, alpha=0.8)
            ax_rmsf.set_xlabel('Residue'); ax_rmsf.set_ylabel('RMSF (Å)')
            ax_rmsf.set_title('Per-Residue RMSF', fontweight='bold')
            ax_rmsf.grid(True, alpha=0.2, axis='y')

        # ── Panel D: RMSF distribution ──
        ax_rmsf_dist = fig.add_subplot(gs_[1, 2])
        if self.rmsf_data:
            ax_rmsf_dist.hist(self.rmsf_data['values'], bins=30,
                              color=PALETTE[1], alpha=0.8, edgecolor='white')
            ax_rmsf_dist.set_xlabel('RMSF (Å)'); ax_rmsf_dist.set_ylabel('Count')
            ax_rmsf_dist.set_title('RMSF Distribution', fontweight='bold')
            ax_rmsf_dist.grid(True, alpha=0.2)

        # ── Panel E: 2D FEL ──
        ax_fel = fig.add_subplot(gs_[2, :2])
        cv1_label, cv1 = self._cv1()
        cv2_label, cv2 = self._cv2()
        if cv1 is not None and cv2 is not None:
            n = min(len(cv1), len(cv2))
            if n >= 20:
                X, Y, G = compute_fel_2d(cv1[:n], cv2[:n], bins=50, kT=self.kT)
                cf = ax_fel.contourf(X, Y, np.clip(G, 0, 5), levels=15,
                                     cmap='RdYlBu_r', vmin=0, vmax=5, extend='max')
                plt.colorbar(cf, ax=ax_fel, label='ΔG (kcal/mol)')
                ax_fel.set_xlabel(cv1_label); ax_fel.set_ylabel(cv2_label)
                ax_fel.set_title('Free Energy Landscape', fontweight='bold')

        # ── Panel F: contact count or hotspot summary ──
        ax_extra = fig.add_subplot(gs_[2, 2])
        if self.contacts_ts.size > 0:
            ax_extra.plot(self.contacts_ts, color=PALETTE[2], lw=1.0, alpha=0.6)
            ax_extra.axhline(self.contacts_ts.mean(), color='red', ls='--',
                             lw=1.8, label=f'Mean={self.contacts_ts.mean():.1f}')
            ax_extra.set_xlabel('Frame'); ax_extra.set_ylabel('# Contacts')
            ax_extra.set_title('Interface Contacts', fontweight='bold')
            ax_extra.legend(fontsize=8)
            ax_extra.grid(True, alpha=0.2)
        elif self.ppi_result:
            score_A = self.ppi_result.get('score_A', {})
            if score_A:
                top10 = sorted(score_A, key=lambda r: -score_A[r])[:10]
                freqs = [score_A[r] for r in top10]
                labels_bar = [r.split(':')[1] for r in top10]
                ax_extra.barh(range(len(top10)), freqs[::-1],
                              color=PALETTE[0], alpha=0.8)
                ax_extra.set_yticks(range(len(top10)))
                ax_extra.set_yticklabels(labels_bar[::-1], fontsize=8)
                ax_extra.axvline(HOTSPOT_FREQ, color='red', ls='--', lw=1.5)
                ax_extra.set_xlabel('Max contact freq')
                ax_extra.set_title('Top Interface Residues (A)', fontweight='bold')
                ax_extra.grid(True, alpha=0.2, axis='x')

        plt.tight_layout()
        self._savefig(fig, '00_Dashboard.png', dpi=150)

    # ── CV helpers ────────────────────────────────────────────────────────────

    def _cv1(self):
        """Primary CV: backbone RMSD."""
        lbl, arr = self._primary_rmsd()
        if arr.size:
            return lbl or 'RMSD (Å)', arr
        return None, None

    def _cv2(self):
        """Secondary CV: sidechain RMSD, or Lig_wrt_Protein, or contact count."""
        # Sidechain RMSD from eaf
        for k in ('Side chains', 'Sidechain', 'All_Heavy'):
            if k in self.rmsd_data and self.rmsd_data[k].size:
                return k + ' RMSD (Å)', self.rmsd_data[k]
        # Ligand RMSD from dat
        for k in ('Lig_wrt_Protein', 'Prot_Sidechain'):
            if k in self.dat_rmsd and self.dat_rmsd[k].size:
                return k + ' (Å)', self.dat_rmsd[k]
        # Contact count
        if self.contacts_ts.size > 1:
            return '# Interface Contacts', self.contacts_ts
        return None, None

    # ── Plot 08 – Binding Site Shape ─────────────────────────────────────────

    def plot_binding_site_shape(self, receptor_pdb: str, ligand_pdb: str = None):
        """
        Generate a binding site shape figure using 3D Cα coordinates from PDB
        file(s).  Requires PPI contact data from a prior EAF parse.

        Panels
        ------
        A  2D projection of full backbone (interface-plane PCA view) with
           binding footprint convex hull and contact-frequency colour coding.
        B  Side view (90° rotation) showing pocket depth.
        C  Interaction network graph: nodes = interface residues,
           edges = inter-chain contacts coloured by interaction type.
        D  Binding-pocket residue-property composition (pie chart).
        """
        if not self.ppi_result:
            print("  No PPI data – cannot draw binding site shape.")
            return

        # ── 1. Parse PDB(s) ──────────────────────────────────────────────────
        rec = _parse_pdb_ca(receptor_pdb)
        lig = _parse_pdb_ca(ligand_pdb) if ligand_pdb else {}

        if not rec:
            print(f"  No Cα atoms found in {receptor_pdb}")
            return

        # Filter ligand to residues near the receptor interface centroid
        # (avoids cluttering the plot with a long full-length CAPON chain)
        if lig:
            # Compute centroid of receptor Cα
            all_rec_coords = np.array([v['coords'] for v in rec.values()])
            rec_centre = all_rec_coords.mean(axis=0)
            lig_coords_all = np.array([v['coords'] for v in lig.values()])
            dists = np.linalg.norm(lig_coords_all - rec_centre, axis=1)
            # Keep only the closest 30 residues or those within 30 Å of receptor centre
            cutoff = max(30.0, np.percentile(dists, 30))
            keep_mask = dists <= cutoff
            lig_items = list(lig.items())
            lig = {k: v for (k, v), keep in zip(lig_items, keep_mask) if keep}

        # ── 2. Map contact scores onto receptor residues ──────────────────────
        score_A  = self.ppi_result.get('score_A',  {})
        hotspot_A = set(self.ppi_result.get('hotspot_A', []))
        iface_A   = set(self.ppi_result.get('iface_A',  []))

        def _freq(chain, resid):
            key = f"{chain}:{rec[(chain, resid)]['resname']}_{resid}" \
                  if (chain, resid) in rec else None
            # score_A keys look like 'A:GLY_1'
            for k, v in score_A.items():
                kc, rest = k.split(':')
                krid = int(rest.split('_')[1])
                if kc == chain and krid == resid:
                    return v
            return 0.0

        # Build arrays: position, freq, label
        coords_rec, labels_rec, freqs_rec = [], [], []
        for (ch, rid), info in sorted(rec.items(), key=lambda x: x[0]):
            coords_rec.append(info['coords'])
            freqs_rec.append(_freq(ch, rid))
            labels_rec.append(f"{ch}:{info['resname']}_{rid}")
        coords_rec = np.array(coords_rec)
        freqs_rec  = np.array(freqs_rec)

        # Ligand backbone (CAPON peptide)
        coords_lig = np.array([v['coords'] for v in lig.values()]) if lig else np.empty((0, 3))
        labels_lig = [f"{ch}:{info['resname']}_{rid}"
                      for (ch, rid), info in lig.items()] if lig else []

        # ── 3. PCA to find interface-optimal viewing plane ────────────────────
        # Use interface residue coordinates for PCA
        iface_mask = freqs_rec >= IFACE_FREQ
        if iface_mask.sum() < 3:
            iface_mask = freqs_rec > 0
        if iface_mask.sum() < 3:
            iface_mask = np.ones(len(freqs_rec), dtype=bool)

        iface_coords = coords_rec[iface_mask]
        pca = _simple_pca(iface_coords)           # returns (n, 3) projected coords
        # pca[:,0] = PC1 (largest spread), pca[:,1] = PC2, pca[:,2] = PC3 (normal)

        # Project all receptor atoms onto the same PC axes
        centre = iface_coords.mean(axis=0)
        coords_c = coords_rec - centre
        pca_vecs = _pca_components(iface_coords)   # shape (3, 3) — rows are PCs
        proj_rec  = coords_c @ pca_vecs.T          # (n_res, 3)
        proj_lig  = ((coords_lig - centre) @ pca_vecs.T) if coords_lig.size else np.empty((0, 3))

        # ── 4. Colour mapping ─────────────────────────────────────────────────
        def _node_color(freq):
            if freq >= HOTSPOT_FREQ:
                return '#D7191C'
            if freq >= IFACE_FREQ:
                return '#FDAE61'
            if freq > 0:
                return '#ABD9E9'
            return '#DDDDDD'

        node_colors = [_node_color(f) for f in freqs_rec]
        node_sizes  = [max(20, f * 300 + 20) for f in freqs_rec]

        # ── 5. Convex hull of hotspot + interface residues ─────────────────────
        from scipy.spatial import ConvexHull
        hull_mask = freqs_rec >= IFACE_FREQ
        hull_pts  = proj_rec[hull_mask, :2]

        # ── 6. Build figure ───────────────────────────────────────────────────
        fig = plt.figure(figsize=(18, 14))
        fig.suptitle(f'Binding Site Shape – {self.label}',
                     fontsize=16, fontweight='bold')
        gs_ = gridspec.GridSpec(2, 2, figure=fig, hspace=0.35, wspace=0.30)

        chains = self.ppi_result.get('all_chains', ['A', 'B'])

        # ── Panel A: Top view (PC1 vs PC2) ───────────────────────────────────
        ax_top = fig.add_subplot(gs_[0, 0])
        _draw_backbone_2d(ax_top, proj_rec[:, 0], proj_rec[:, 1],
                          freqs_rec, labels_rec, node_colors, node_sizes,
                          hull_pts, chains[0])
        if proj_lig.size:
            ax_top.scatter(proj_lig[:, 0], proj_lig[:, 1],
                           marker='^', s=120, c='#31A354', zorder=6,
                           edgecolors='darkgreen', linewidths=1.2,
                           label=f'Chain {chains[1] if len(chains) > 1 else "B"} (ligand)')
            # Draw ligand backbone line
            ax_top.plot(proj_lig[:, 0], proj_lig[:, 1],
                        '-', color='#31A354', lw=1.5, alpha=0.7, zorder=5)
        ax_top.set_xlabel('PC1 (Å)', fontsize=11)
        ax_top.set_ylabel('PC2 (Å)', fontsize=11)
        ax_top.set_title(f'Interface Plane View\n(PC1 vs PC2)', fontsize=12,
                         fontweight='bold')
        ax_top.set_aspect('equal')
        ax_top.legend(fontsize=8, loc='upper right', framealpha=0.8)

        # ── Panel B: Side view (PC1 vs PC3) ──────────────────────────────────
        ax_side = fig.add_subplot(gs_[0, 1])
        _draw_backbone_2d(ax_side, proj_rec[:, 0], proj_rec[:, 2],
                          freqs_rec, labels_rec, node_colors, node_sizes,
                          None, chains[0], label_hotspots=False)
        if proj_lig.size:
            ax_side.scatter(proj_lig[:, 0], proj_lig[:, 2],
                            marker='^', s=120, c='#31A354', zorder=6,
                            edgecolors='darkgreen', linewidths=1.2)
            ax_side.plot(proj_lig[:, 0], proj_lig[:, 2],
                         '-', color='#31A354', lw=1.5, alpha=0.7, zorder=5)

        # Pocket depth annotation
        if hull_mask.sum() > 0:
            depth = proj_rec[hull_mask, 2].max() - proj_rec[hull_mask, 2].min()
            ax_side.axhspan(proj_rec[hull_mask, 2].min(),
                            proj_rec[hull_mask, 2].max(),
                            alpha=0.08, color='gold',
                            label=f'Pocket depth ≈ {depth:.1f} Å')
            ax_side.legend(fontsize=9)

        ax_side.set_xlabel('PC1 (Å)', fontsize=11)
        ax_side.set_ylabel('PC3 / depth (Å)', fontsize=11)
        ax_side.set_title('Side View\n(interface depth)', fontsize=12, fontweight='bold')
        ax_side.set_aspect('equal')

        # Legend patches for both projection panels
        legend_patches = [
            Patch(color='#D7191C', label=f'Hotspot (≥{HOTSPOT_FREQ:.0%})'),
            Patch(color='#FDAE61', label=f'Interface (≥{IFACE_FREQ:.0%})'),
            Patch(color='#ABD9E9', label='Peripheral'),
            Patch(color='#DDDDDD', label='Non-contact'),
        ]
        ax_top.legend(handles=legend_patches + ([
            plt.Line2D([0], [0], marker='^', color='w', markerfacecolor='#31A354',
                       markersize=9, label='Ligand Cα')
        ] if proj_lig.size else []), fontsize=8, framealpha=0.85, loc='best')

        # ── Panel C: Interaction network ──────────────────────────────────────
        ax_net = fig.add_subplot(gs_[1, 0])
        _draw_interaction_network(ax_net, self.ppi_result, proj_rec, labels_rec)
        ax_net.set_title('Interface Interaction Network', fontsize=12, fontweight='bold')

        # ── Panel D: Binding pocket residue properties ────────────────────────
        ax_pie = fig.add_subplot(gs_[1, 1])
        _draw_pocket_composition(ax_pie, rec, labels_rec, freqs_rec, IFACE_FREQ)
        ax_pie.set_title('Binding Pocket Residue Properties', fontsize=12,
                         fontweight='bold')

        plt.tight_layout()
        self._savefig(fig, '08_BindingSiteShape.png')

    # ── Plot 09 – RMSD vs Rg FEL ─────────────────────────────────────────────

    def plot_rmsd_rg_fel(self):
        """3-panel: Rg time series | RMSD time series | 2D FEL (RMSD vs Rg)."""
        _, rmsd_arr = self._primary_rmsd()
        rg = self.rg_data
        if rmsd_arr.size == 0:
            print("  No RMSD data for RMSD–Rg FEL – skipping.")
            return
        if rg.size == 0:
            print("  No Rg data for RMSD–Rg FEL – skipping.")
            return

        n = min(len(rmsd_arr), len(rg))
        if n < 20:
            return
        rmsd_arr = rmsd_arr[:n]
        rg       = rg[:n]
        frames   = np.arange(n)

        fig, axes = plt.subplots(1, 3, figsize=(17, 5))
        ax_rg, ax_rmsd, ax_fel = axes

        # Rg time series
        ax_rg.plot(frames, rg, color=PALETTE[2], lw=0.8, alpha=0.45)
        avg_rg = self._running_avg(rg, max(5, n // 50))
        ax_rg.plot(frames, avg_rg, color=PALETTE[2], lw=2.2,
                   label=f'Mean = {rg.mean():.2f} Å')
        ax_rg.set_xlabel('Frame', fontsize=12)
        ax_rg.set_ylabel('Radius of Gyration (Å)', fontsize=12)
        ax_rg.set_title('Rg Time Series', fontweight='bold', fontsize=13)
        ax_rg.legend(fontsize=9)
        ax_rg.grid(True, alpha=0.25)

        # RMSD time series
        ax_rmsd.plot(frames, rmsd_arr, color=PALETTE[0], lw=0.8, alpha=0.45)
        avg_rmsd = self._running_avg(rmsd_arr, max(5, n // 50))
        ax_rmsd.plot(frames, avg_rmsd, color=PALETTE[0], lw=2.2,
                     label=f'Mean = {rmsd_arr.mean():.2f} Å')
        ax_rmsd.set_xlabel('Frame', fontsize=12)
        ax_rmsd.set_ylabel('RMSD (Å)', fontsize=12)
        ax_rmsd.set_title('RMSD Time Series', fontweight='bold', fontsize=13)
        ax_rmsd.legend(fontsize=9)
        ax_rmsd.grid(True, alpha=0.25)

        # 2D FEL
        X, Y, G = compute_fel_2d(rmsd_arr, rg, bins=50, sigma=1.5, kT=self.kT)
        G_plot  = np.clip(G, 0, 5.0)
        cf = ax_fel.contourf(X, Y, G_plot, levels=15, cmap='RdYlBu_r',
                             vmin=0, vmax=5, extend='max')
        ax_fel.contour(X, Y, G_plot, levels=8, colors='black', alpha=0.3, linewidths=0.5)
        cb = plt.colorbar(cf, ax=ax_fel, pad=0.02)
        cb.set_label('ΔG (kcal/mol)', fontsize=10)
        min_idx = np.unravel_index(G_plot.argmin(), G_plot.shape)
        ax_fel.plot(X[min_idx], Y[min_idx], 'w*', markersize=16, zorder=5,
                    label=f'Min ({X[min_idx]:.2f}, {Y[min_idx]:.2f})')
        ax_fel.set_xlabel('RMSD (Å)', fontsize=12)
        ax_fel.set_ylabel('Rg (Å)', fontsize=12)
        ax_fel.set_title('RMSD vs Rg FEL', fontweight='bold', fontsize=13)
        ax_fel.legend(fontsize=8)

        plt.suptitle(f'RMSD–Rg Free Energy Landscape – {self.label}',
                     fontsize=14, fontweight='bold')
        plt.tight_layout()
        self._savefig(fig, '09_RMSD_Rg_FEL.png')

    # ── Plot 10 – Secondary Structure Timeline ────────────────────────────────

    def plot_ss_timeline(self):
        """Heatmap of SS codes per residue per frame + per-residue occupancy bar."""
        if self.ss_matrix is None or self.ss_matrix.size == 0:
            print("  No secondary structure data – skipping SS timeline.")
            return

        mat = np.where(self.ss_matrix < 0, 0, self.ss_matrix)   # remap −1 → 0
        n_res, n_frames = mat.shape

        ss_cmap   = matplotlib.colors.ListedColormap(
            ['#EEEEEE', '#E74C3C', '#3498DB', '#F39C12'])
        ss_bounds = [-0.5, 0.5, 1.5, 2.5, 3.5]
        ss_norm   = matplotlib.colors.BoundaryNorm(ss_bounds, ss_cmap.N)
        ss_labels = {0: 'Coil/Loop', 1: 'α-Helix', 2: 'β-Strand', 3: 'Turn'}
        ss_colors_d = {0: '#EEEEEE', 1: '#E74C3C', 2: '#3498DB', 3: '#F39C12'}

        fig = plt.figure(figsize=(16, max(7, n_res * 0.09 + 3)))
        gs_ = gridspec.GridSpec(1, 2, figure=fig, wspace=0.06,
                                width_ratios=[4, 1])
        ax_heat = fig.add_subplot(gs_[0])
        ax_bar  = fig.add_subplot(gs_[1])

        ax_heat.imshow(mat, aspect='auto', origin='lower',
                       cmap=ss_cmap, norm=ss_norm, interpolation='nearest')

        step = max(1, n_res // 25)
        ax_heat.set_yticks(range(0, n_res, step))
        if self.ss_residues:
            ax_heat.set_yticklabels(
                [self.ss_residues[i].split(':')[1] if ':' in self.ss_residues[i]
                 else self.ss_residues[i] for i in range(0, n_res, step)],
                fontsize=7)
        frame_step = max(1, n_frames // 10)
        ax_heat.set_xticks(range(0, n_frames, frame_step))
        ax_heat.set_xticklabels(
            [str(i) for i in range(0, n_frames, frame_step)], fontsize=8)
        ax_heat.set_xlabel('Frame', fontsize=12)
        ax_heat.set_ylabel('Residue', fontsize=12)
        ax_heat.set_title('Secondary Structure Timeline', fontweight='bold', fontsize=13)

        legend_patches = [Patch(color=ss_colors_d[k], label=v)
                          for k, v in ss_labels.items()]
        ax_heat.legend(handles=legend_patches, loc='upper right', fontsize=8,
                       framealpha=0.85, ncol=2)

        # Stacked occupancy bars
        ss_types  = [1, 2, 3, 0]
        occupancy = np.array([[np.mean(mat[i] == t) for t in ss_types]
                               for i in range(n_res)])
        left = np.zeros(n_res)
        for j, t in enumerate(ss_types):
            ax_bar.barh(range(n_res), occupancy[:, j], left=left,
                        color=ss_colors_d[t], alpha=0.9,
                        label=ss_labels[t])
            left += occupancy[:, j]

        ax_bar.set_xlim(0, 1)
        ax_bar.set_xlabel('Occupancy', fontsize=11)
        ax_bar.set_yticks([])
        ax_bar.set_ylim(ax_heat.get_ylim())
        ax_bar.set_title('SS Occupancy', fontweight='bold', fontsize=12)
        ax_bar.grid(True, alpha=0.2, axis='x')
        ax_bar.legend(handles=[Patch(color=ss_colors_d[t], label=ss_labels[t])
                                for t in ss_types],
                      loc='lower right', fontsize=8, framealpha=0.8)

        plt.suptitle(f'Secondary Structure Analysis – {self.label}',
                     fontsize=14, fontweight='bold')
        plt.tight_layout()
        self._savefig(fig, '10_SecondaryStructure_Timeline.png')

    # ── Plot 11 – Contact Persistence Timeline ────────────────────────────────

    def plot_contact_persistence(self):
        """Binary heatmap of per-residue interface contacts over time."""
        if self.contact_persist is None or self.contact_persist.size == 0:
            print("  No contact-persistence data – skipping.")
            return

        mat     = self.contact_persist   # (n_res, n_frames) int8
        n_res, n_frames = mat.shape

        freq  = mat.mean(axis=1)
        order = np.argsort(freq)[::-1]
        mat   = mat[order]
        labels = [self.contact_persist_labels[i] for i in order]
        freq   = freq[order]

        keep  = freq >= 0.05
        if keep.sum() == 0:
            keep = np.ones(len(freq), dtype=bool)
        mat    = mat[keep]
        labels = [l for l, k in zip(labels, keep) if k]
        freq   = freq[keep]
        n_show = len(labels)

        fig = plt.figure(figsize=(16, max(6, n_show * 0.32 + 3)))
        gs_ = gridspec.GridSpec(1, 2, figure=fig, wspace=0.04,
                                width_ratios=[4, 1])
        ax_heat = fig.add_subplot(gs_[0])
        ax_freq = fig.add_subplot(gs_[1])

        ax_heat.imshow(mat, aspect='auto', origin='upper',
                       cmap='Blues', vmin=0, vmax=1, interpolation='nearest')

        step = max(1, n_show // 25)
        ax_heat.set_yticks(range(0, n_show, step))
        ax_heat.set_yticklabels(
            [labels[i].split(':')[1] if ':' in labels[i] else labels[i]
             for i in range(0, n_show, step)], fontsize=7)
        frame_step = max(1, n_frames // 10)
        ax_heat.set_xticks(range(0, n_frames, frame_step))
        ax_heat.set_xticklabels(
            [str(i) for i in range(0, n_frames, frame_step)], fontsize=8)
        ax_heat.set_xlabel('Frame', fontsize=12)
        ax_heat.set_ylabel('Residue (sorted by frequency)', fontsize=12)
        ax_heat.set_title('Contact Persistence Timeline', fontweight='bold', fontsize=13)

        # Star hotspot residues
        if self.ppi_result:
            hotset = set(self.ppi_result.get('hotspot_A', []) +
                         self.ppi_result.get('hotspot_B', []))
            for i, lbl in enumerate(labels):
                if lbl in hotset:
                    ax_heat.annotate('★', xy=(-1, i), xycoords='data',
                                     ha='right', va='center',
                                     fontsize=8, color='gold')

        # Frequency bar
        bar_colors = [
            '#D7191C' if f >= HOTSPOT_FREQ else
            '#FDAE61' if f >= IFACE_FREQ   else '#ABD9E9'
            for f in freq
        ]
        ax_freq.barh(range(n_show), freq, color=bar_colors, alpha=0.9)
        ax_freq.axvline(HOTSPOT_FREQ, color='red',    ls='--', lw=1.5, alpha=0.8)
        ax_freq.axvline(IFACE_FREQ,   color='orange', ls='--', lw=1.5, alpha=0.8)
        ax_freq.set_xlim(0, 1.05)
        ax_freq.set_xlabel('Contact frequency', fontsize=11)
        ax_freq.set_yticks([])
        ax_freq.set_ylim(n_show - 0.5, -0.5)
        ax_freq.set_title('Frequency', fontweight='bold', fontsize=12)
        ax_freq.grid(True, alpha=0.2, axis='x')
        ax_freq.legend(handles=[
            Patch(color='#D7191C', label=f'Hotspot ≥{HOTSPOT_FREQ:.0%}'),
            Patch(color='#FDAE61', label=f'Interface ≥{IFACE_FREQ:.0%}'),
            Patch(color='#ABD9E9', label='Peripheral'),
        ], fontsize=8, loc='lower right', framealpha=0.85)

        plt.suptitle(f'Interface Contact Persistence – {self.label}',
                     fontsize=14, fontweight='bold')
        plt.tight_layout()
        self._savefig(fig, '11_ContactPersistence.png')

    # ── Plot 12 – Differential RMSF ──────────────────────────────────────────

    def plot_differential_rmsf(self):
        """ΔRMSF = RMSF_bound − RMSF_unbound, highlighting allosteric changes."""
        if not self.rmsf_data or not self.unbound_rmsf:
            print("  Need bound + unbound RMSF – skipping differential RMSF.")
            return

        bound_res    = self.rmsf_data['residues']
        bound_vals   = self.rmsf_data['values']
        unbound_map  = {r: v for r, v in zip(self.unbound_rmsf['residues'],
                                              self.unbound_rmsf['values'])}

        aligned_labels, delta, bound_a, unbound_a = [], [], [], []
        for r, b in zip(bound_res, bound_vals):
            if r in unbound_map:
                aligned_labels.append(r)
                delta.append(b - unbound_map[r])
                bound_a.append(b)
                unbound_a.append(unbound_map[r])

        if not aligned_labels:
            print("  No overlapping residues between bound/unbound – skipping.")
            return

        delta      = np.array(delta)
        bound_a    = np.array(bound_a)
        unbound_a  = np.array(unbound_a)

        resids = []
        for r in aligned_labels:
            m = re.match(r'[A-Za-z]:\w+_(\d+)', r)
            resids.append(int(m.group(1)) if m else len(resids))
        resids = np.array(resids)

        iface_set  = set()
        if self.ppi_result:
            iface_set = set(
                self.ppi_result.get('hotspot_A', []) +
                self.ppi_result.get('iface_A',   []))
        iface_mask = np.array([r in iface_set for r in aligned_labels])

        fig, axes = plt.subplots(2, 1, figsize=(14, 9),
                                 gridspec_kw={'height_ratios': [2, 1.2]})
        ax_d, ax_abs = axes

        # ΔRMSF bar chart
        bar_c = np.where(delta > 0, '#D7191C', '#2C7BB6')
        ax_d.bar(resids, delta, color=bar_c, alpha=0.82, width=0.9)
        ax_d.axhline(0, color='black', lw=1.0)
        thr_hi = delta.mean() + 2 * delta.std()
        thr_lo = delta.mean() - 2 * delta.std()
        ax_d.axhline(thr_hi, color='red',  ls='--', lw=1.5, alpha=0.75,
                     label=f'+2σ = {thr_hi:+.2f} Å')
        ax_d.axhline(thr_lo, color='blue', ls='--', lw=1.5, alpha=0.75,
                     label=f'−2σ = {thr_lo:+.2f} Å')

        for rid, lbl, d_val, is_iface in zip(resids, aligned_labels,
                                              delta, iface_mask):
            if is_iface:
                offset = 6 if d_val >= 0 else -14
                ax_d.annotate('◆', xy=(rid, d_val),
                              xytext=(0, offset), textcoords='offset points',
                              fontsize=8, color='purple', ha='center')

        ax_d.set_xlabel('Residue Number', fontsize=12)
        ax_d.set_ylabel('ΔRMSF  bound − unbound (Å)', fontsize=12)
        ax_d.set_title('Differential RMSF – Allosteric Effect Map',
                        fontweight='bold', fontsize=13)
        legend_h = [
            Patch(color='#D7191C', label='More flexible when bound'),
            Patch(color='#2C7BB6', label='Less flexible when bound'),
            plt.Line2D([0], [0], color='red',  ls='--', label=f'+2σ = {thr_hi:+.2f} Å'),
            plt.Line2D([0], [0], color='blue', ls='--', label=f'−2σ = {thr_lo:+.2f} Å'),
        ]
        if iface_mask.any():
            legend_h.append(plt.Line2D(
                [0], [0], marker='D', color='w', markerfacecolor='purple',
                markersize=9, label='Interface residue'))
        ax_d.legend(handles=legend_h, fontsize=9, framealpha=0.85)
        ax_d.grid(True, alpha=0.2, axis='y')

        # Absolute RMSF overlay
        ax_abs.fill_between(resids, bound_a,   alpha=0.25, color=PALETTE[0])
        ax_abs.plot(resids, bound_a,   color=PALETTE[0], lw=1.8,
                    label=f'Bound  (μ={bound_a.mean():.2f} Å)')
        ax_abs.fill_between(resids, unbound_a, alpha=0.25, color=PALETTE[1])
        ax_abs.plot(resids, unbound_a, color=PALETTE[1], lw=1.8, ls='--',
                    label=f'Unbound (μ={unbound_a.mean():.2f} Å)')

        for rid, is_iface in zip(resids, iface_mask):
            if is_iface:
                ax_abs.axvline(rid, color='purple', lw=0.5, alpha=0.35)

        ax_abs.set_xlabel('Residue Number', fontsize=12)
        ax_abs.set_ylabel('RMSF (Å)', fontsize=12)
        ax_abs.set_title('Absolute RMSF – Bound vs Unbound', fontweight='bold', fontsize=12)
        ax_abs.legend(fontsize=9, framealpha=0.85)
        ax_abs.grid(True, alpha=0.2, axis='y')

        plt.suptitle(f'Differential RMSF Analysis – {self.label}',
                     fontsize=14, fontweight='bold')
        plt.tight_layout()
        self._savefig(fig, '12_Differential_RMSF.png')

    # ── Run all ───────────────────────────────────────────────────────────────

    def run(self, receptor_pdb: str = None, ligand_pdb: str = None,
            unbound_eaf: str = None):
        print('\n' + '=' * 60)
        print('  SCHRÖDINGER DESMOND MD ANALYSIS')
        print('=' * 60)
        self.load()
        if unbound_eaf:
            self.load_unbound_rmsf(unbound_eaf)
        print('\n── Generating figures ──')
        self.plot_rmsd()
        self.plot_rmsf()
        self.plot_fel_2d()
        self.plot_fel_3d()
        self.plot_ppi_contact_map()
        self.plot_ppi_hotspots()
        self.plot_interaction_breakdown()
        if receptor_pdb and os.path.isfile(receptor_pdb):
            self.plot_binding_site_shape(receptor_pdb, ligand_pdb)
        self.plot_dashboard()
        self.plot_rmsd_rg_fel()
        self.plot_ss_timeline()
        self.plot_contact_persistence()
        self.plot_differential_rmsf()
        self._print_summary()
        print(f'\nAll figures saved to: {self.output_dir}/')

    def _print_summary(self):
        print('\n' + '─' * 60)
        print('SUMMARY')
        print('─' * 60)
        lbl, arr = self._primary_rmsd()
        if arr.size:
            print(f"RMSD ({lbl}): {arr.mean():.2f} ± {arr.std():.2f} Å  "
                  f"[{arr.min():.2f}–{arr.max():.2f}]")
        if self.rmsf_data and 'values' in self.rmsf_data:
            vals = self.rmsf_data['values']
            res  = self.rmsf_data['residues']
            top3 = np.argsort(vals)[-3:][::-1]
            top3_str = ', '.join(f"{res[i]} ({vals[i]:.2f} Å)" for i in top3)
            print(f"Most flexible residues: {top3_str}")
        if self.ppi_result:
            print(f"Inter-chain contact pairs: {self.ppi_result.get('n_inter_pairs', 0)}")
            print(f"Chain A hotspots (≥{HOTSPOT_FREQ:.0%}): "
                  f"{[r.split(':')[1] for r in self.ppi_result.get('hotspot_A', [])]}")
            print(f"Chain A interface (≥{IFACE_FREQ:.0%}): "
                  f"{[r.split(':')[1] for r in self.ppi_result.get('iface_A', [])]}")
            print(f"Chain B hotspots (≥{HOTSPOT_FREQ:.0%}): "
                  f"{[r.split(':')[1] for r in self.ppi_result.get('hotspot_B', [])]}")
            print(f"Chain B interface (≥{IFACE_FREQ:.0%}): "
                  f"{[r.split(':')[1] for r in self.ppi_result.get('iface_B', [])]}")


# ══════════════════════════════════════════════════════════════════════════════
# PDB HELPERS
# ══════════════════════════════════════════════════════════════════════════════

def _parse_pdb_ca(filepath: str) -> dict:
    """
    Parse a PDB file and return Cα atoms.
    Returns: {(chain, resid): {'resname': str, 'coords': np.array([x,y,z])}}
    """
    atoms = {}
    with open(filepath, errors='replace') as fh:
        for line in fh:
            rec = line[:6].strip()
            if rec not in ('ATOM', 'HETATM'):
                continue
            name = line[12:16].strip()
            if name != 'CA':
                continue
            chain  = line[21]
            try:
                resid  = int(line[22:26])
                x, y, z = float(line[30:38]), float(line[38:46]), float(line[46:54])
            except ValueError:
                continue
            resname = line[17:20].strip()
            key = (chain, resid)
            if key not in atoms:   # keep first occurrence (alt-loc A)
                atoms[key] = {'resname': resname, 'coords': np.array([x, y, z])}
    return atoms


def _pca_components(coords: np.ndarray) -> np.ndarray:
    """
    Return the top-3 PCA eigenvectors for coords as rows of a (3×3) matrix.
    Falls back to identity if too few points.
    """
    if coords.shape[0] < 3:
        return np.eye(3)
    cen = coords.mean(axis=0)
    cov = np.cov((coords - cen).T)
    vals, vecs = np.linalg.eigh(cov)
    # Sort descending
    order = np.argsort(vals)[::-1]
    return vecs[:, order].T   # rows are PC1, PC2, PC3


def _simple_pca(coords: np.ndarray) -> np.ndarray:
    """Project coords onto top-3 PCs. Returns (n, 3)."""
    pcs = _pca_components(coords)
    cen = coords.mean(axis=0)
    return (coords - cen) @ pcs.T


def _draw_backbone_2d(ax, x, y, freqs, labels, colors, sizes,
                      hull_pts, chain_label, label_hotspots=True):
    """Draw a Cα backbone trace on ax with coloured nodes."""
    # Backbone line
    ax.plot(x, y, '-', color='#CCCCCC', lw=0.8, alpha=0.6, zorder=1)

    # Scatter
    sc = ax.scatter(x, y, c=colors, s=sizes, zorder=3,
                    edgecolors='white', linewidths=0.5)

    # Convex hull of interface residues
    if hull_pts is not None and len(hull_pts) >= 3:
        from scipy.spatial import ConvexHull
        try:
            hull = ConvexHull(hull_pts)
            poly_x = np.append(hull_pts[hull.vertices, 0],
                               hull_pts[hull.vertices[0], 0])
            poly_y = np.append(hull_pts[hull.vertices, 1],
                               hull_pts[hull.vertices[1], 1])
            ax.fill(poly_x, poly_y, alpha=0.10, color='gold', zorder=2)
            ax.plot(poly_x, poly_y, '--', color='goldenrod', lw=1.8,
                    alpha=0.8, zorder=2, label='Binding footprint')
            # Centroid
            cx, cy = hull_pts[:, 0].mean(), hull_pts[:, 1].mean()
            ax.plot(cx, cy, 'k+', markersize=14, markeredgewidth=2,
                    zorder=5, label='Site centroid')
        except Exception:
            pass

    # Label hotspot residues
    if label_hotspots:
        for xi, yi, lbl, freq in zip(x, y, labels, freqs):
            if freq >= HOTSPOT_FREQ:
                short = lbl.split(':')[1] if ':' in lbl else lbl
                ax.annotate(short, (xi, yi), textcoords='offset points',
                            xytext=(4, 4), fontsize=7, color='#8B0000',
                            fontweight='bold', zorder=6)

    ax.set_facecolor('#F8F8F8')
    ax.grid(True, alpha=0.15)


def _draw_interaction_network(ax, ppi_result: dict,
                              proj_rec: np.ndarray, labels_rec: list):
    """
    Draw the inter-chain interaction network using approximate 2D positions
    derived from the PCA projection of receptor Cα coordinates.
    Nodes = interface residues (both chains).
    Edges = inter-chain contacts from ProtProtInter, coloured by type.
    """
    inter_pairs = ppi_result.get('inter_pairs', {})
    score_A     = ppi_result.get('score_A', {})
    score_B     = ppi_result.get('score_B', {})
    chains      = ppi_result.get('all_chains', ['A', 'B'])

    if not inter_pairs:
        ax.text(0.5, 0.5, 'No inter-chain contacts found',
                ha='center', va='center', transform=ax.transAxes)
        return

    # ── Node positions ────────────────────────────────────────────────────────
    # Chain A nodes: use PC1/PC2 of their Cα
    label_to_proj: dict = {}
    for i, lbl in enumerate(labels_rec):
        label_to_proj[lbl] = proj_rec[i, :2]

    # Chain B nodes: arrange in a horizontal line above the A cluster
    b_nodes = sorted(set(ppi_result.get('hotspot_B', []) +
                         ppi_result.get('iface_B', [])),
                     key=_resid_sort_key)
    if not b_nodes:
        b_nodes = sorted({rB for (rA, rB) in inter_pairs}, key=_resid_sort_key)

    # Position chain B above the binding site centroid
    a_iface_proj = np.array([label_to_proj[r] for r in score_A
                             if r in label_to_proj and score_A[r] >= IFACE_FREQ])
    if a_iface_proj.size:
        cx, cy = a_iface_proj.mean(axis=0)
        y_top = a_iface_proj[:, 1].max() + 8
    else:
        cx, cy, y_top = 0.0, 0.0, 20.0

    spread = max(10.0, len(b_nodes) * 4.0)
    for i, r in enumerate(b_nodes):
        bx = cx - spread / 2 + i * spread / max(1, len(b_nodes) - 1)
        label_to_proj[r] = np.array([bx, y_top])

    # ── Draw edges ────────────────────────────────────────────────────────────
    itype_drawn = set()
    for (rA, rB), itypes in inter_pairs.items():
        if rA not in label_to_proj or rB not in label_to_proj:
            continue
        pA = label_to_proj[rA]
        pB = label_to_proj[rB]
        dominant = max(itypes, key=itypes.get)
        freq     = max(itypes.values())
        color    = ITYPE_COLORS.get(dominant, '#888888')
        lw       = 1.0 + freq * 3.0
        lbl      = ITYPE_LABELS.get(dominant, dominant) if dominant not in itype_drawn else None
        ax.plot([pA[0], pB[0]], [pA[1], pB[1]],
                '-', color=color, lw=lw, alpha=0.65, zorder=1,
                label=lbl)
        if lbl:
            itype_drawn.add(dominant)

    # ── Draw nodes ────────────────────────────────────────────────────────────
    def _plot_nodes(node_list, score_dict, node_color, marker, zorder=3):
        for r in node_list:
            if r not in label_to_proj:
                continue
            px, py = label_to_proj[r]
            freq   = score_dict.get(r, 0)
            size   = 80 + freq * 300
            c      = '#D7191C' if freq >= HOTSPOT_FREQ else '#FDAE61'
            ax.scatter(px, py, c=c, s=size, marker=marker, zorder=zorder,
                       edgecolors='white', linewidths=0.8)
            short = r.split(':')[1] if ':' in r else r
            ax.annotate(short, (px, py), textcoords='offset points',
                        xytext=(3, 3), fontsize=7, fontweight='bold', zorder=zorder + 1)

    all_A = sorted(set(rA for (rA, rB) in inter_pairs), key=_resid_sort_key)
    all_B = sorted(set(rB for (rA, rB) in inter_pairs), key=_resid_sort_key)
    _plot_nodes(all_A, score_A, PALETTE[0], 'o', zorder=4)
    _plot_nodes(all_B, score_B, PALETTE[1], '^', zorder=4)

    # Chain labels
    if a_iface_proj.size:
        ax.text(a_iface_proj[:, 0].mean(), a_iface_proj[:, 1].min() - 6,
                f'Chain {chains[0]} (receptor)', ha='center', fontsize=9,
                color=PALETTE[0], fontweight='bold')
    ax.text(cx, y_top + 6, f'Chain {chains[1] if len(chains) > 1 else "B"} (ligand)',
            ha='center', fontsize=9, color=PALETTE[1], fontweight='bold')

    ax.legend(fontsize=8, loc='lower right', framealpha=0.8, ncol=2)
    ax.set_facecolor('#F8F8F8')
    ax.grid(True, alpha=0.12)
    ax.set_xlabel('PC1 (Å)', fontsize=10)
    ax.set_ylabel('PC2 (Å)', fontsize=10)


# Amino acid property classification
_AA_PROPS = {
    'ARG': 'Pos. charged', 'LYS': 'Pos. charged', 'HIS': 'Pos. charged',
    'ASP': 'Neg. charged', 'GLU': 'Neg. charged',
    'SER': 'Polar',  'THR': 'Polar',  'ASN': 'Polar',  'GLN': 'Polar',
    'TYR': 'Polar',  'CYS': 'Polar',
    'GLY': 'Hydrophobic', 'ALA': 'Hydrophobic', 'VAL': 'Hydrophobic',
    'ILE': 'Hydrophobic', 'LEU': 'Hydrophobic', 'MET': 'Hydrophobic',
    'PHE': 'Hydrophobic', 'TRP': 'Hydrophobic', 'PRO': 'Hydrophobic',
}
_PROP_COLORS = {
    'Pos. charged': '#3182BD',
    'Neg. charged': '#E6550D',
    'Polar':        '#74C476',
    'Hydrophobic':  '#9ECAE1',
}


def _draw_pocket_composition(ax, rec: dict, labels: list,
                             freqs: np.ndarray, iface_thresh: float):
    """Pie chart of residue physicochemical properties at the binding interface."""
    prop_counts: dict = {}
    for lbl, freq in zip(labels, freqs):
        if freq < iface_thresh:
            continue
        resname = lbl.split(':')[1].split('_')[0] if ':' in lbl else ''
        prop = _AA_PROPS.get(resname, 'Other')
        prop_counts[prop] = prop_counts.get(prop, 0) + 1

    if not prop_counts:
        ax.text(0.5, 0.5, 'No interface residues found',
                ha='center', va='center', transform=ax.transAxes)
        return

    labels_pie = list(prop_counts.keys())
    sizes_pie  = [prop_counts[p] for p in labels_pie]
    colors_pie = [_PROP_COLORS.get(p, '#CCCCCC') for p in labels_pie]

    wedges, texts, autotexts = ax.pie(
        sizes_pie, labels=labels_pie, colors=colors_pie,
        autopct='%1.0f%%', pctdistance=0.75, startangle=90,
        wedgeprops={'edgecolor': 'white', 'linewidth': 2})
    for t in texts:
        t.set_fontsize(10)
    for at in autotexts:
        at.set_fontsize(9)

    # Annotate with actual residue lists
    for prop in labels_pie:
        members = [lbl.split(':')[1] for lbl, f in zip(labels, freqs)
                   if f >= iface_thresh and
                   _AA_PROPS.get(lbl.split(':')[1].split('_')[0], 'Other') == prop]
        if members:
            ax.annotate(', '.join(members),
                        xy=(0, -1.45 - 0.18 * labels_pie.index(prop)),
                        fontsize=7.5, ha='center',
                        color=_PROP_COLORS.get(prop, '#555'))

    ax.set_title('')   # title set by caller


# ══════════════════════════════════════════════════════════════════════════════
# UTILITIES
# ══════════════════════════════════════════════════════════════════════════════

def _resid_sort_key(res: str) -> tuple:
    """Sort residue labels 'A:GLY_10' by chain then number."""
    m = re.match(r'([A-Za-z]):\w+_(\d+)', res)
    if m:
        return (m.group(1), int(m.group(2)))
    return ('', 0)


def _lighten(hex_color: str, amount: float = 0.5) -> str:
    """Lighten a hex colour by blending with white."""
    hex_color = hex_color.lstrip('#')
    r, g, b = (int(hex_color[i:i+2], 16) for i in (0, 2, 4))
    r = int(r + (255 - r) * amount)
    g = int(g + (255 - g) * amount)
    b = int(b + (255 - b) * amount)
    return f'#{r:02x}{g:02x}{b:02x}'


def auto_discover(path: str) -> tuple:
    """
    Given a path (directory, .eaf file, or job name prefix), return
    (eaf_files, dat_dir).
    """
    eaf_files, dat_dir = [], None

    if os.path.isfile(path) and path.endswith('.eaf'):
        eaf_files = [path]
        parent = os.path.dirname(path)
        # Look for raw-data/ sibling
        for cand in ('raw-data', 'rawdata', 'data'):
            d = os.path.join(parent, cand)
            if os.path.isdir(d):
                dat_dir = d
                break
    elif os.path.isdir(path):
        eaf_files = sorted(glob.glob(os.path.join(path, '*.eaf')))
        for cand in ('raw-data', 'rawdata', 'data'):
            d = os.path.join(path, cand)
            if os.path.isdir(d):
                dat_dir = d
                break

    return eaf_files, dat_dir


# ══════════════════════════════════════════════════════════════════════════════
# CLI
# ══════════════════════════════════════════════════════════════════════════════

def build_parser() -> argparse.ArgumentParser:
    p = argparse.ArgumentParser(
        prog='desmond_analysis.py',
        description=textwrap.dedent("""\
            Schrödinger Desmond MD Analysis Tool
            Generates RMSD, RMSF, 2D/3D FEL, and PPI binding-site figures
            from .eaf and .dat output files."""),
        formatter_class=argparse.RawDescriptionHelpFormatter,
        epilog=textwrap.dedent("""\
            Examples
            --------
            # Single simulation (EAF + DAT):
              python desmond_analysis.py \\
                -eaf "schrodinger work files/desmond_md_job_CAPON_bound_NOS-out_pl_1.eaf" \\
                -dat "schrodinger work files/CAPON-bound-nNOS-MD/raw-data/" \\
                -o capon_analysis/ -T 300 --label nNOS-CAPON

            # Auto-discover from a directory:
              python desmond_analysis.py --auto "schrodinger work files/CAPON-bound-nNOS-MD/"

            # Multiple replicate EAF files:
              python desmond_analysis.py -eaf rep1.eaf rep2.eaf rep3.eaf -o combined/
            """)
    )
    p.add_argument('-eaf', '--eaf', nargs='+', metavar='FILE',
                   help='.eaf file(s) (Schrödinger Event Analysis Files)')
    p.add_argument('-dat', '--dat', metavar='DIR',
                   help='Directory containing .dat files (raw-data/)')
    p.add_argument('-o', '--output', default='desmond_analysis_output',
                   help='Output directory (default: desmond_analysis_output)')
    p.add_argument('-T', '--temperature', type=float, default=300.0,
                   help='Simulation temperature in K (default: 300)')
    p.add_argument('--label', default=None,
                   help='Label for figure titles (default: derived from filename)')
    p.add_argument('--auto', metavar='PATH',
                   help='Auto-discover .eaf and .dat files under PATH')
    p.add_argument('-rec', '--receptor-pdb', metavar='FILE',
                   help='Receptor .pdb file for binding-site shape figure')
    p.add_argument('-lig', '--ligand-pdb', metavar='FILE',
                   help='Ligand/peptide .pdb file for binding-site shape figure')
    p.add_argument('--hotspot-cutoff', type=float, default=0.70,
                   help='Contact frequency threshold for hotspot (default: 0.70)')
    p.add_argument('--iface-cutoff', type=float, default=0.30,
                   help='Contact frequency threshold for interface (default: 0.30)')
    p.add_argument('--unbound-eaf', metavar='FILE',
                   help='Unbound simulation .eaf file for differential RMSF analysis')
    return p


def main():
    parser = build_parser()
    args = parser.parse_args()

    # Apply thresholds
    global HOTSPOT_FREQ, IFACE_FREQ
    HOTSPOT_FREQ = args.hotspot_cutoff
    IFACE_FREQ   = args.iface_cutoff

    eaf_files, dat_dir = [], args.dat

    # Auto-discover
    if args.auto:
        eaf_files, discovered_dat = auto_discover(args.auto)
        if dat_dir is None:
            dat_dir = discovered_dat
        if not eaf_files:
            sys.exit(f"ERROR: No .eaf files found under {args.auto}")
    elif args.eaf:
        eaf_files = args.eaf
    else:
        parser.print_help()
        sys.exit('\nERROR: Provide -eaf FILE(s) or --auto PATH')

    # Validate
    missing = [f for f in eaf_files if not os.path.isfile(f)]
    if missing:
        sys.exit(f"ERROR: File(s) not found: {missing}")

    analysis = DesmondAnalysis(
        eaf_files   = eaf_files,
        dat_dir     = dat_dir,
        output_dir  = args.output,
        temperature = args.temperature,
        label       = args.label,
    )
    analysis.run(
        receptor_pdb=args.receptor_pdb,
        ligand_pdb=args.ligand_pdb,
        unbound_eaf=args.unbound_eaf,
    )


if __name__ == '__main__':
    main()
```
